# A statistical framework for powerful multi-trait rare variant analysis in large-scale whole-genome sequencing studies

**DOI:** 10.1101/2023.10.30.564764

**Authors:** Xihao Li, Han Chen, Margaret Sunitha Selvaraj, Eric Van Buren, Hufeng Zhou, Yuxuan Wang, Ryan Sun, Zachary R. McCaw, Zhi Yu, Donna K. Arnett, Joshua C. Bis, John Blangero, Eric Boerwinkle, Donald W. Bowden, Jennifer A. Brody, Brian E. Cade, April P. Carson, Jenna C. Carlson, Nathalie Chami, Yii-Der Ida Chen, Joanne E. Curran, Paul S. de Vries, Myriam Fornage, Nora Franceschini, Barry I. Freedman, Charles Gu, Nancy L. Heard-Costa, Jiang He, Lifang Hou, Yi-Jen Hung, Marguerite R. Irvin, Robert C. Kaplan, Sharon L.R. Kardia, Tanika Kelly, Iain Konigsberg, Charles Kooperberg, Brian G. Kral, Changwei Li, Ruth J.F. Loos, Michael C. Mahaney, Lisa W. Martin, Rasika A. Mathias, Ryan L. Minster, Braxton D. Mitchell, May E. Montasser, Alanna C. Morrison, Nicholette D. Palmer, Patricia A. Peyser, Bruce M. Psaty, Laura M. Raffield, Susan Redline, Alexander P. Reiner, Stephen S. Rich, Colleen M. Sitlani, Jennifer A. Smith, Kent D. Taylor, Hemant Tiwari, Ramachandran S. Vasan, Zhe Wang, Lisa R. Yanek, Bing Yu, NHLBI Trans-Omics for Precision Medicine (TOPMed) Consortium, Kenneth M. Rice, Jerome I. Rotter, Gina M. Peloso, Pradeep Natarajan, Zilin Li, Zhonghua Liu, Xihong Lin

**Affiliations:** Department of Biostatistics, University of North Carolina at Chapel Hill, Chapel Hill, NC, USA; Department of Genetics, University of North Carolina at Chapel Hill, Chapel Hill, NC, USA; Human Genetics Center, Department of Epidemiology, Human Genetics, and Environmental Sciences, School of Public Health, The University of Texas Health Science Center at Houston, Houston, TX, USA; Center for Precision Health, School of Biomedical Informatics, The University of Texas Health Science Center at Houston, Houston, TX, USA; Center for Genomic Medicine and Cardiovascular Research Center, Massachusetts General Hospital, Boston, MA, USA; Program in Medical and Population Genetics, Broad Institute of Harvard and MIT, Cambridge, MA, USA; Department of Medicine, Harvard Medical School, Boston, MA, USA; Department of Biostatistics, Harvard T.H. Chan School of Public Health, Boston, MA, USA; Department of Biostatistics, Boston University School of Public Health, Boston, MA, USA; Department of Biostatistics, University of Texas MD Anderson Cancer Center, Houston, TX, USA; Provost Office, University of South Carolina, Columbia, SC, USA; Cardiovascular Health Research Unit, Department of Medicine, University of Washington, Seattle, WA, USA; Department of Human Genetics and South Texas Diabetes and Obesity Institute, School of Medicine, The University of Texas Rio Grande Valley, Brownsville, TX, USA; Human Genome Sequencing Center, Baylor College of Medicine, Houston, TX, USA; Department of Biochemistry, Wake Forest University School of Medicine, Winston-Salem, NC, USA; Division of Sleep and Circadian Disorders, Brigham and Women’s Hospital, Boston, MA, USA; Division of Sleep Medicine, Harvard Medical School, Boston, MA, USA; Department of Medicine, University of Mississippi Medical Center, Jackson, MS, USA; Department of Human Genetics and Department of Biostatistics, University of Pittsburgh, Pittsburgh, PA, USA; The Charles Bronfman Institute for Personalized Medicine, Icahn School of Medicine at Mount Sinai, New York, NY, USA; The Institute for Translational Genomics and Population Sciences, Department of Pediatrics, The Lundquist Institute for Biomedical Innovation at Harbor-UCLA Medical Center, Torrance, CA, USA; Brown Foundation Institute of Molecular Medicine, McGovern Medical School, the University of Texas Health Science Center at Houston, Houston, TX, USA; Department of Epidemiology, University of North Carolina at Chapel Hill, Chapel Hill, NC, USA; Department of Internal Medicine, Nephrology, Wake Forest University School of Medicine, Winston-Salem, NC, USA; Division of Biology & Biomedical Sciences, Washington University School of Medicine, St. Louis, MO, USA; Department of Neurology, Boston University Chobanian & Avedisian School of Medicine, Boston, MA, USA; Framingham Heart Study, Framingham, MA, USA; Department of Epidemiology, Tulane University School of Public Health and Tropical Medicine, New Orleans, LA, USA; Tulane University Translational Science Institute, New Orleans, LA, USA; Department of Preventive Medicine, Northwestern University, Chicago, IL, USA; Department of Internal Medicine, Tri-Service General Hospital, National Defense Medical Center, Taipei, Taiwan; Department of Epidemiology, School of Public Health, University of Alabama at Birmingham, Birmingham, AL, USA; Department of Epidemiology and Population Health, Albert Einstein College of Medicine, Bronx, NY, USA; Division of Public Health Sciences, Fred Hutchinson Cancer Center, Seattle, WA, USA; Department of Epidemiology, School of Public Health, University of Michigan, Ann Arbor, MI, USA; Department of Medicine, Division of Nephrology, University of Illinois Chicago, Chicago, IL, USA; Department of Biomedical Informatics, University of Colorado, Aurora, CO, USA; Department of Medicine, Johns Hopkins University School of Medicine, Baltimore, MD, USA; Novo Nordisk Foundation Center for Basic Metabolic Research, Faculty of Health and Medical Sciences, University of Copenhagen, Copenhagen, Denmark; George Washington University School of Medicine and Health Sciences, Washington, DC, USA; Department of Medicine, University of Maryland School of Medicine, Baltimore, MD, USA; Departments of Epidemiology, University of Washington, Seattle, WA, USA; Department of Health Systems and Population Health, University of Washington, Seattle, WA, USA; Center for Public Health Genomics, University of Virginia, Charlottesville, VA, USA; Department of Biostatistics, School of Public Health, University of Alabama at Birmingham, Birmingham, AL, USA; Department of Quantitative and Qualitative Health Sciences, UT Health San Antonio School of Public Health, San Antonia, TX, USA; Department of Biostatistics, University of Washington, Seattle, WA, USA; Department of Biostatistics, Mailman School of Public Health, Columbia University, New York, NY, USA; Department of Statistics, Harvard University, Cambridge, MA, USA

## Abstract

Large-scale whole-genome sequencing (WGS) studies have improved our understanding of the contributions of coding and noncoding rare variants to complex human traits. Leveraging association effect sizes across multiple traits in WGS rare variant association analysis can improve statistical power over single-trait analysis, and also detect pleiotropic genes and regions. Existing multi-trait methods have limited ability to perform rare variant analysis of large-scale WGS data. We propose MultiSTAAR, a statistical framework and computationally-scalable analytical pipeline for functionally-informed multi-trait rare variant analysis in large-scale WGS studies. MultiSTAAR accounts for relatedness, population structure and correlation among phenotypes by jointly analyzing multiple traits, and further empowers rare variant association analysis by incorporating multiple functional annotations. We applied MultiSTAAR to jointly analyze three lipid traits (low-density lipoprotein cholesterol, high-density lipoprotein cholesterol and triglycerides) in 61,861 multi-ethnic samples from the Trans-Omics for Precision Medicine (TOPMed) Program. We discovered new associations with lipid traits missed by single-trait analysis, including rare variants within an enhancer of *NIPSNAP3A* and an intergenic region on chromosome 1.

---

Advances in next generation sequencing technologies and the decreasing cost of whole-exome/whole-genome sequencing (WES/WGS) have made it possible to study the genetic underpinnings of rare variants (i.e. minor allele frequency (MAF) < 1%) in complex human traits. Large nationwide consortia and biobanks, such as the National Heart, Lung and Blood Institute (NHLBI)’s Trans-Omics for Precision Medicine (TOPMed) Program^1^, the National Human Genome Research Institute’s Genome Sequencing Program (GSP), the National Institute of Health’s All of Us Research Program^2^, and the UK’s Biobank WGS Program^3^, are expected to sequence more than a million of individuals in total, at more than 1 billion genetic variants in both coding and noncoding regions of the human genome, while also recording thousands of phenotypes. To mitigate the lack of power of single-variant analyses to identify rare variant associations^4^, variant set tests have been proposed to analyze the joint effects of multiple rare variants ^5-9^, where most of the work has focused single trait analysis.

Pleiotropy occurs when genetic variants influence multiple traits^10^. There is growing empirical evidence from genome-wide association studies (GWASs) that many variants have pleiotropic effects^11,12^. Identifying these effects can provide valuable insights into the genetic architecture of complex traits^13^. As such, it is of increasing interest to identify pleiotropic rare variants by jointly analyzing multiple traits in WGS rare variant association studies (RVASs).

Several existing methods for multi-trait rare variant association analysis, such as MSKAT^14^, Multi-SKAT^15^ and MTAR^16^, have shown that leveraging the cross-phenotype correlation structure can improve the power of multi-trait analyses compared to single-trait analyses when analyzing pleiotropic genes^14-17^. However, existing methods do not scale well, and are not feasible when analyzing large-scale WGS studies with hundreds of millions of rare variants in samples exhibiting relatedness and population structure.

Furthermore, none of the existing multi-trait rare variant analysis methods leverages functional annotations that predict the biological functionality of variants, resulting in limited interpretability and power loss. While the STAAR method^18^ dynamically incorporates multiple variant functional annotations to maximize the power of rare variant association tests, it is designed for single-trait analysis and cannot be directly applied to multiple traits.

To overcome these limitations, we propose the Multi-trait variant-Set Test for Association using Annotation infoRmation (MultiSTAAR), a statistical framework for multi-trait rare variant analyses of large-scale WGS studies and biobanks. It has several features. First, by fitting a null Multivariate Linear Mixed Model (MLMM)^19^ for multiple quantitative traits simultaneously, adjusting for ancestry principal components (PCs)^20^ and using a sparse genetic relatedness matrix (GRM)^21,22^, MultiSTAAR scales well but also accounts for relatedness and population structure, as well as correlations among the multiple traits. Second, MultiSTAAR enables the incorporation of multiple variant functional annotations as weights to improve the power of RVASs. Furthermore, we provide MultiSTAAR via a comprehensive pipeline for large-scale WGS studies, that facilitates functionally-informed multi-trait analysis of both coding and noncoding rare variants. Third, MultiSTAAR enables conditional multi-trait analysis to assess rare variant association signals beyond known common and low frequency variants.

In the current study, we conducted extensive simulation studies to demonstrate the validity of MultiSTAAR and to assess the power gain of MultiSTAAR by incorporating multiple relevant variant functional annotations, and its ability in preserving Type I error rates. We then applied MultiSTAAR to perform WGS RVAS of 61,838 ancestrally diverse participants from 20 studies from NHLBI’s TOPMed consortium by jointly analyzing three circulating lipid traits: low-density lipoprotein cholesterol (LDL-C), high-density lipoprotein cholesterol (HDL-C) and triglycerides (TG). We show that MultiSTAAR is computationally feasible for large-scale WGS multi-trait rare variant analysis, and in conditional analysis of LDL-C, HDL-C and TG, MultiSTAAR identifies signals that were missed either by the existing multi-trait rare variant analysis methods that overlook variant functional annotations, or by single-trait functionally-informed analysis that ignore correlations between phenotypes.

## Results

### Overview of the methods

MultiSTAAR is a statistical framework and an analytic pipeline for jointly analyzing multiple traits in large-scale WGS rare variant association studies. There are two main components in the MultiSTAAR framework: (i) fitting null MLMMs using ancestry PCs and sparse GRMs to account for population structure, relatedness and the correlation between phenotypes, and (ii) testing for associations between each aggregated variant set and multiple traits by dynamically incorporating multiple variant functional annotations^18^ (**Fig. 1a**).

**Fig. 1.**
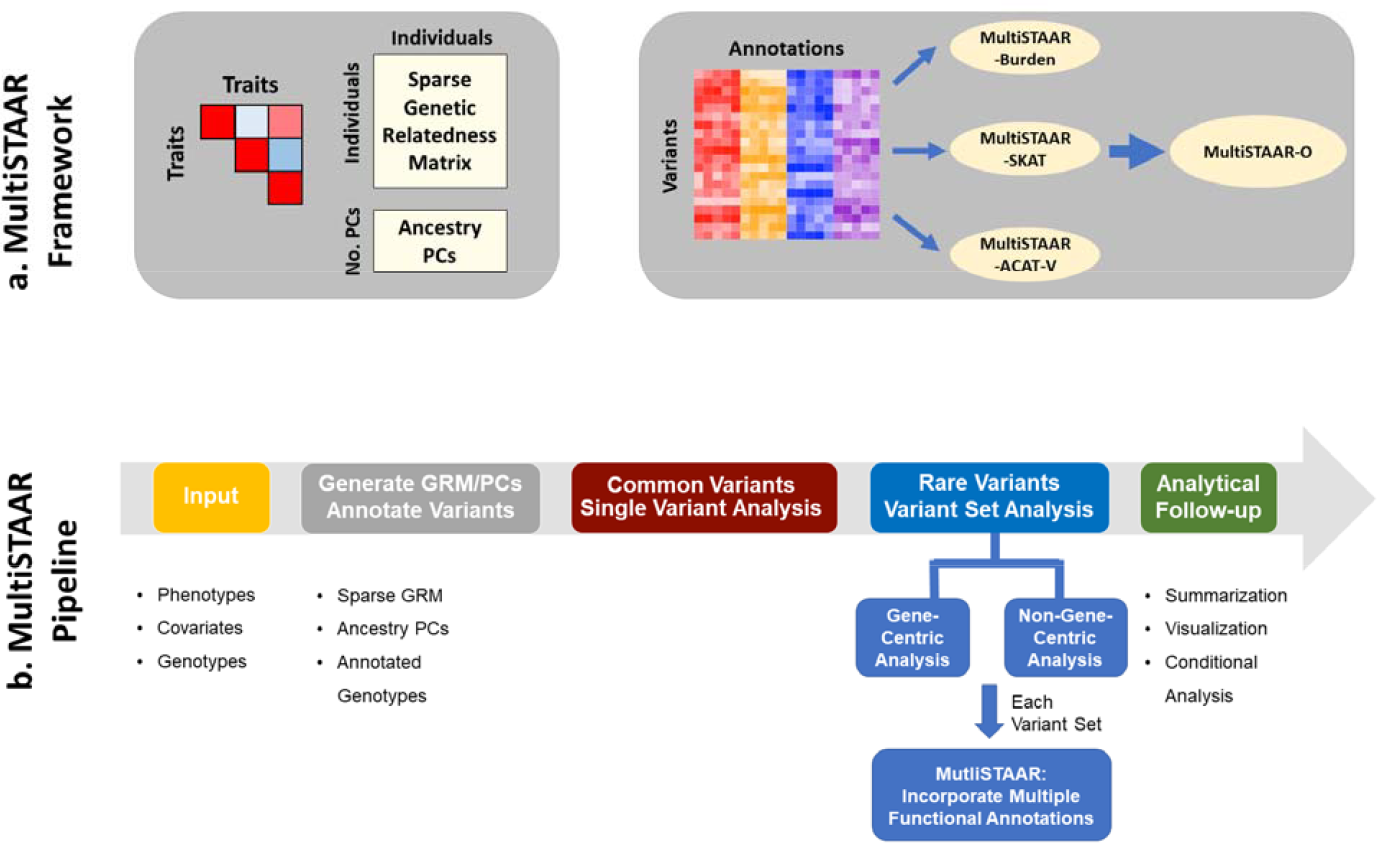
MultiSTAAR framework and pipeline. **a**, MultiSTAAR framework. (i) Fit null Multivariate Linear Mixed Models (MLMMs) using sparse GRM and ancestry PCs to account for population structure, relatedness and the correlation between phenotypes. (ii) Test for associations between each variant set and multiple traits by dynamically incorporating multiple variant functional annotations. **b**, MultiSTAAR pipeline. (i) Prepare the input data of MultiSTAAR, including genotypes, multiple phenotypes and covariates. (ii) Calculate sparse GRM, ancestry PCs and annotate all variants in the genome. (iii) Perform single variant analysis for common variants. (iv) Define the rare variant analysis units, including gene-centric analysis of five coding functional categories and eight noncoding functional categories and non-gene-centric analysis of sliding windows. (v) Provide result summarization and perform analytical follow-up via conditional analysis.

In WGS RVASs, an important but often underemphasized challenge is selecting biologically-meaningful and functionally-interpretable analysis units, especially for the noncoding genome^23,24^. In gene-centric analyses of multiple traits, MultiSTAAR provides five functional categories (masks) to aggregate coding rare variants of each protein-coding gene, as well as an additional eight masks of regulatory regions to aggregate noncoding rare variants. In non-gene-centric analyses of multiple traits, MultiSTAAR performs agnostic genetic region analyses using sliding windows^18,25^ (**Fig. 1b**).

For each rare variant set analyzed, MultiSTAAR first constructs the multi-trait burden, SKAT and ACAT-V test statistics (**Methods**). For each type of rare variant test, MultiSTAAR calculates multiple candidate *P* values using different variant functional annotations as weights, following the STAAR framework^18^. MultiSTAAR then aggregates the association strength by combining the *P* values from all annotations using the ACAT method, that provides robustness to correlation between tests^9^, and proposes an omnibus test, MultiSTAAR-O, that leverages the advantages of different type of tests (**Methods**). Furthermore, MultiSTAAR can test multi-trait rare variants’ associations conditional on a set of known associations (**Fig. 1b**).

### Simulation studies

To evaluate the type I error rates and the power of MultiSTAAR, we performed simulation studies under several configurations. Following the steps described in Data Simulation (**Methods**), we generated three quantitative traits with a correlation matrix similar to the empirical correlation in the three lipid traits^26-28^. We then generated genotypes by simulating 20,000 sequences for 100 different 1 megabase (Mb) regions, each of them were generated to mimic the linkage disequilibrium structure of an African American population by using the calibration coalescent model^29^. Throughout the simulation studies, we randomly and uniformly selected 5-kilobase (kb) regions from these 1-Mb regions and considered sample sizes of 10,000 for each replicate. The simulation studies focused on aggregating uncommon variants with an MAF < 5%.

### Type I error rate evaluations

We performed 10^8^ simulations to evaluate the type I error rates of the multi-trait burden, SKAT, ACAT-V and MultiSTAAR-O tests at *α=*10^−4^, 10^−5^ and 10^−6^ (**Supplementary Table 1**). The results show that, for multi-trait rare variant analysis, all four MultiSTAAR tests controlled the type I error rates at very close to the nominal *α*levels.

### Empirical power simulations

We next assessed the power of MultiSTAAR-O for the analysis of multiple phenotypes under different genetic architectures, while also comparing its power with existing methods. Specifically, we considered four models, in which variants in the signal region (variant-phenotype association regions) were associated with (1) one phenotype only, (2) two positively correlated phenotypes, (3) two negatively correlated phenotypes and (4) all three phenotypes. In addition, we considered different proportions (5%, 15% and 35% on average) of causal variants in the signal region, where causality of variants depended on different sets of annotations, and the effect size directions of causal variants were allowed to vary (**Methods**). Power was evaluated as the proportions of *P* values less than *α=*10^−7^ based on 10^4^ simulations. Overall, MultiSTAAR-O consistently delivered higher power to detect signal regions compared to multi-trait burden, SKAT and ACAT-V tests, through its incorporation of multiple annotations (**Extended Data Figs. 2-5, Supplementary Figs. 1-4**). This power advantage was also robust to the existence of noninformative annotations.

### Application to the TOPMed lipids WGS data

We applied MultiSTAAR to identify rare variant associations with three quantitative lipid traits (LDL-C, HDL-C and TG) through a multi-trait analysis using TOPMed Freeze 8 WGS data, comprising 61,838 individuals from 20 multi-ethnic studies (**Supplementary Note**). LDL-C values were adjusted for the usage of lipid-lowering medication^26,30^ (**Methods**), and DNA samples were sequenced at >30x target coverage. Sample- and variant-level quality control were performed for each participating study^1,26,30^.

Race/ethnicity was measured using a combination of self-reported race/ethnicity and study recruitment information^31^ (**Supplementary Note**). Of the 61,838 samples, 15,636 (25.3%) were Black or African American, 27,439 (44.4%) were White, 4,461 (7.2%) were Asian or Asian American, 13,138 (21.2%) were Hispanic/Latino American and 1,164 (1.9%) were Samoans. There were 414 million single-nucleotide variants (SNVs) observed overall, with 6.5 million (1.6%) common variants (MAF > 5%), 5.2 million (1.2%) low-frequency variants (1% ≤ MAF ≤ 5%) and 402 million (97.2%) rare variants (MAF < 1%). The study-specific demographics and baseline characteristics are given in **Supplementary Table 2**.

### Gene-centric multi-trait analysis of coding and noncoding rare variants

We applied MultiSTAAR-O on gene-centric multi-trait analysis of coding and noncoding rare variants of genes with lipid traits in TOPMed. For coding variants, rare variants (MAF < 1%) from five coding functional categories (masks) were aggregated, separately, and analyzed using a joint model for LDL-C, HDL-C and TG, including (1) putative loss-of-function (stop gain, stop loss and splice) rare variants, (2) missense rare variants, (3) disruptive missense rare variants, (4) putative loss-of-function and disruptive missense rare variants and (5) synonymous rare variants of each protein-coding gene. The putative loss-of-function, missense and synonymous RVs were defined by GENCODE Variant Effect Predictor (VEP) categories^32^. The disruptive variants were further defined by MetaSVM^33^, which measures the deleteriousness of missense mutations. We incorporated 9 annotation principal components (aPCs)^18,26,34^, CADD^35^, LINSIGHT^36^, FATHMM-XF^37^ and MetaSVM^33^ (for missense rare variants only) along with the two MAF-based weights^4^ in MultiSTAAR-O (**Supplementary Table 3**). The overall distribution of MultiSTAAR-O *P* values was well-calibrated for the multi-trait analysis of coding rare variants (**Extended Data Fig. 1b**). At a Bonferroni-corrected significance threshold of *α* = 0.05/(20,000 × 5) = 5.00 × 10^−7^, accounting for five different coding masks across protein-coding genes, MultiSTAAR-O identified 51 genome-wide significant associations using unconditional multi-trait analysis (**Extended Data Fig. 1a, Supplementary Table 4**). After conditioning on previously reported variants associated with LDL-C, HDL-C or TG located within a 1 Mb broader region of each coding mask in the GWAS Catalog and Million Veteran Program (MVP)^26,38,39^, 34 out of the 51 associations remained significant at the Bonferroni-corrected threshold of *α=*0.05/51 =9.80 ×10^−4^ (**Table 1**).

**Table 1.**
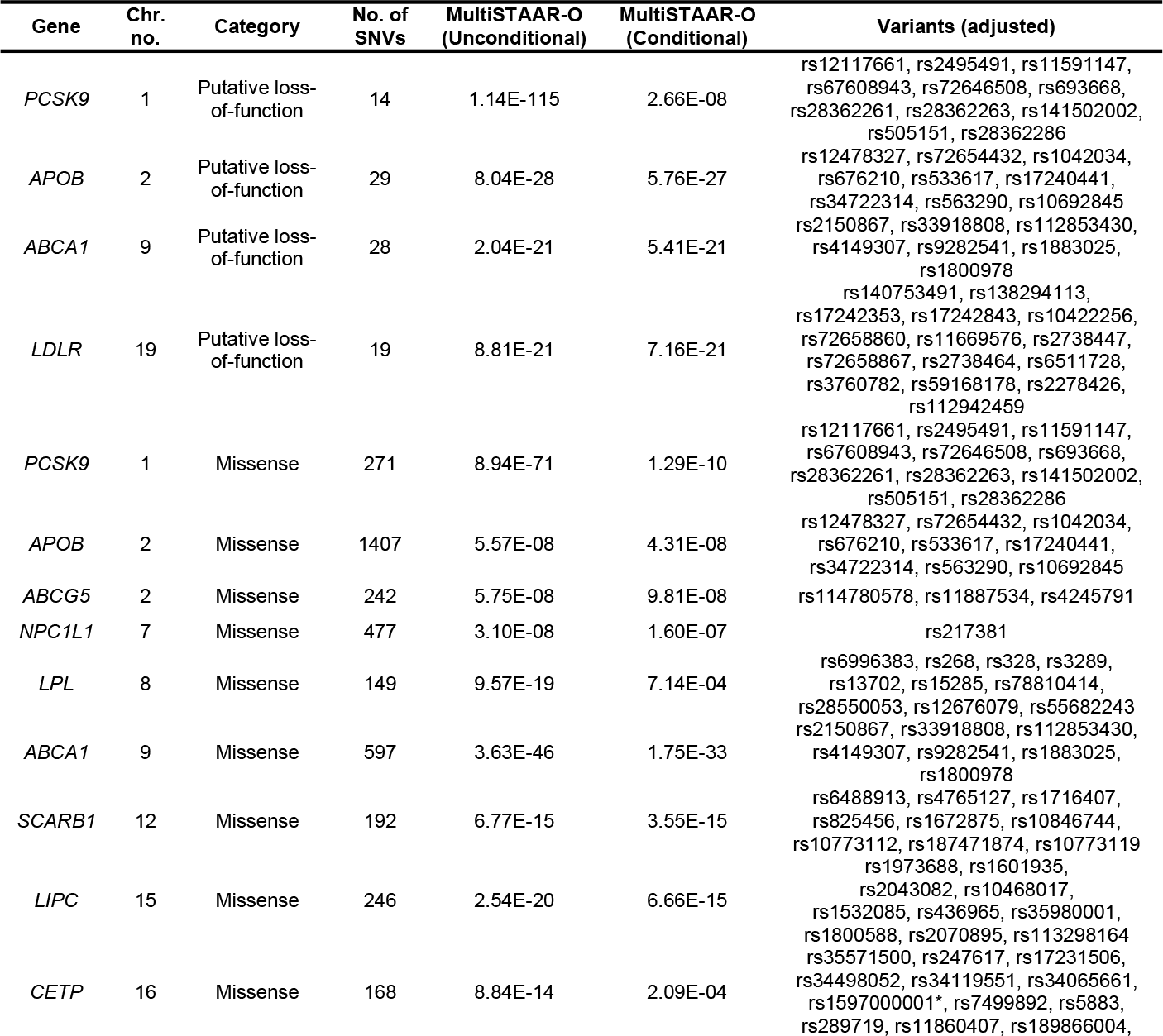
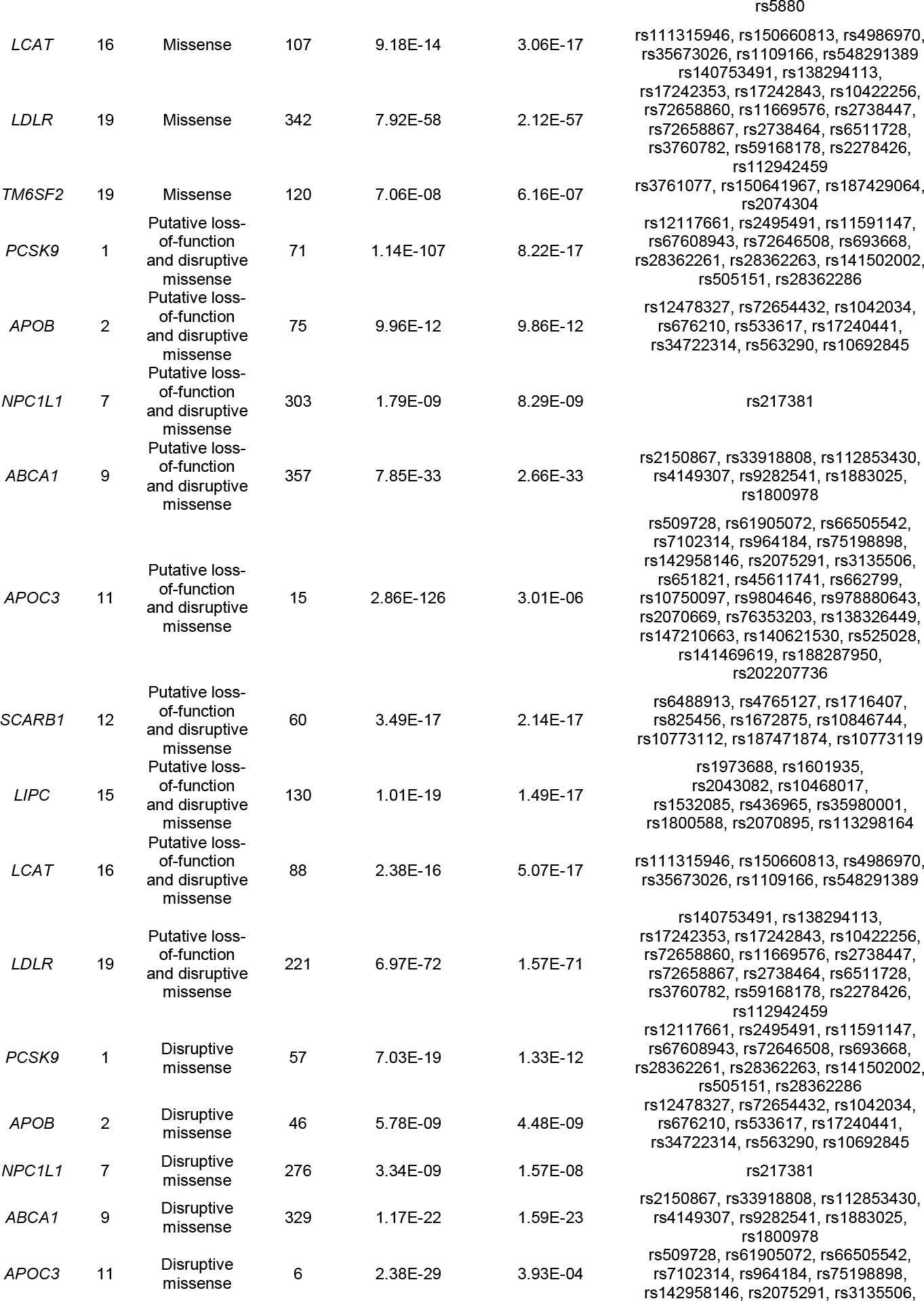
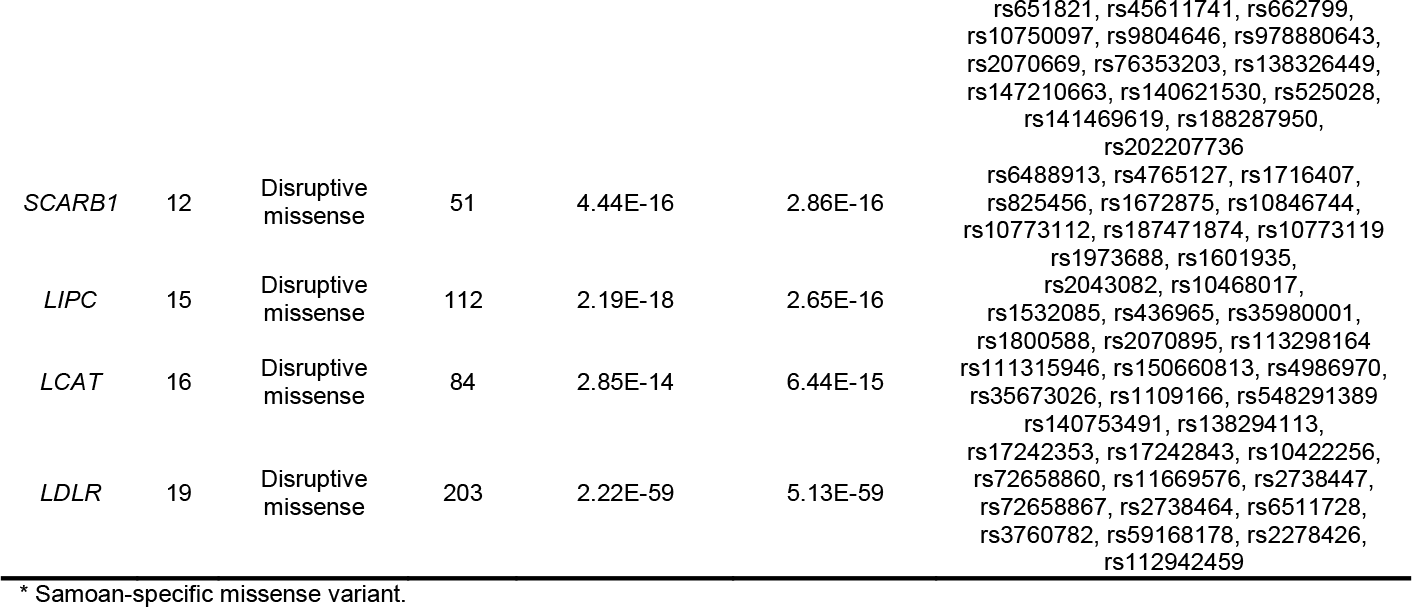
TOPMed Gene-centric coding multi-trait analysis results of both unconditional analysis and analysis conditional on known lipids-associated variants. A total of 61,838 samples from the TOPMed Program were considered in the analysis. Results for the conditionally significant genes (unconditional MultiSTAAR-O *P* < 5.00 × 10^−7^; conditional MultiSTAAR-O *P* < 9.80 × 10^−4^) are presented in the table. MultiSTAAR-O is a two-sided test. Chr. no., chromosome number; Category, functional category; No. of SNVs, number of rare variants (MAF < 1%) of the particular coding functional category in the gene; MultiSTAAR-O, MultiSTAAR-O *P* value; Variants (adjusted), adjusted variants in the conditional analysis.

For non-coding variants, rare variants from eight noncoding masks were analyzed in a similar fashion, including (1) promoter rare variants overlaid with CAGE sites^40^, (2) promoter rare variants overlaid with DHS sites^41^, (3) enhancer rare variants overlaid with CAGE sites^42,43^, (4) enhancer rare variants overlaid with DHS sites^41,43^, (5) untranslated region (UTR) rare variants, (6) upstream region rare variants, (7) downstream region rare variants of each protein-coding gene and (8) rare variants in ncRNA genes^24^. The promoter rare variants were defined as rare variants in the ±3-kilobase (kb) window of transcription start sites with the overlap of CAGE sites or DHS sites. The enhancer rare variants were defined as RVs in GeneHancer-predicted regions with the overlap of CAGE sites or DHS sites. The UTR, upstream, downstream and ncRNA rare variants were defined by GENCODE VEP categories^32^. With a well-calibrated overall distribution of MultiSTAAR-O *P* values (**Extended Data Fig. 1d**) and at a Bonferroni-corrected significance threshold of *α=*0.05/(20,000× 7) =3.57 ×10^−7^, accounting for seven different noncoding masks across protein-coding genes, MultiSTAAR-O identified 76 genome-wide significant associations using unconditional multi-trait analysis (**Extended Data Fig. 1c, Supplementary Table 5**). After conditioning on known lipids-associated variants^26,38,39^, 6 out of the 76 associations remained significant at the Bonferroni-corrected threshold of *α=*0.05/76 =6.58 ×10^−4^ (**Table 2**). These included promoter CAGE and enhancer CAGE rare variants in *APOA1*, promoter DHS rare variants in *CETP*, enhancer CAGE rare variants in *SPC24*, and enhancer DHS rare variants in *NIPSNAP3A* and *LIPC*.

**Table 2.**
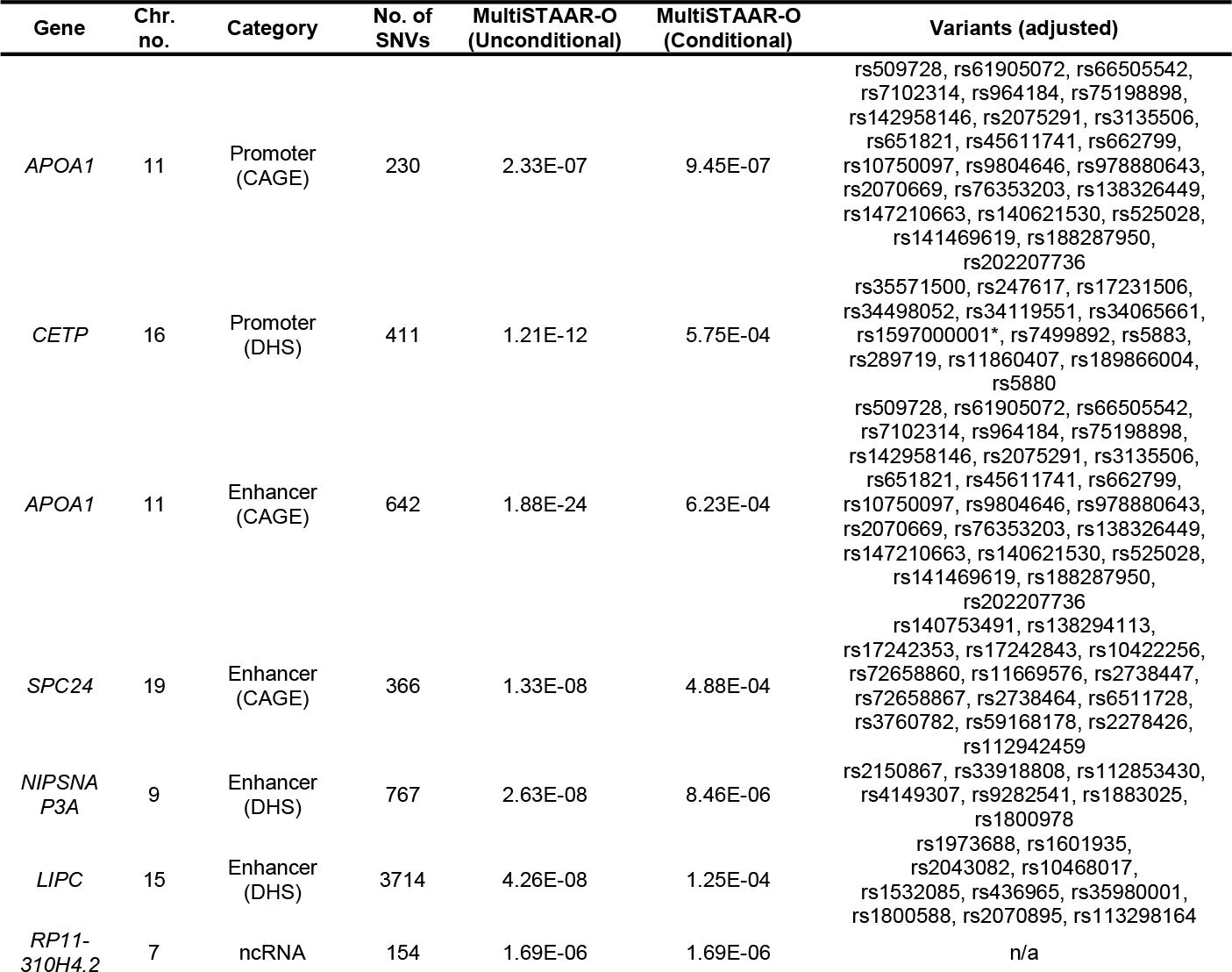
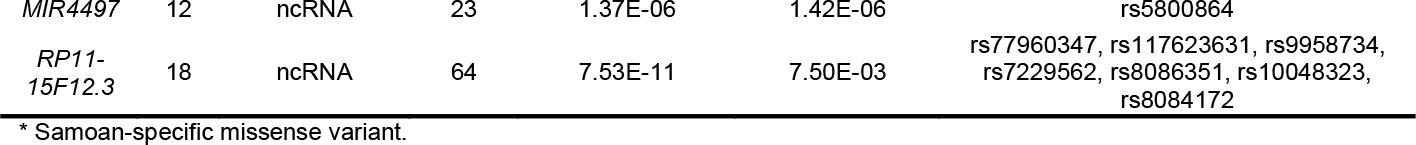
TOPMed Gene-centric noncoding multi-trait analysis results of both unconditional analysis and analysis conditional on known lipids-associated variants. A total of 61,838 samples from the TOPMed Program were considered in the analysis. Results for the conditionally significant genes (unconditional MultiSTAAR-O *P* < 3.57 × 10^−7^ and conditional MultiSTAAR-O *P* < 6.58 × 10^−4^for 7 different noncoding masks across protein-coding genes; unconditional MultiSTAAR-O *P* < 2.50 × 10^−6^ and conditional MultiSTAAR-O *P* < 8.33 × 10^−3^ ncRNA genes) are presented in the table. MultiSTAAR-O is a two-sided test. Chr. no., chromosome number; Category, functional category; No. of SNVs, number of rare variants (MAF < 1%) of the particular noncoding functional category in the gene; MultiSTAAR-O, MultiSTAAR-O *P* value; Variants (adjusted), adjusted variants in the conditional analysis; n/a, no variant adjusted in the conditional analysis.

| Gene | Chr. no. | Category | No. of SNVs | MultiSTAAR-O (Unconditional) | MultiSTAAR-O (Conditional) | Variants (adjusted) |
| --- | --- | --- | --- | --- | --- | --- |
| *APOA1* | 11 | Promoter (CAGE) | 230 | 2.33E-07 | 9.45E-07 | rs509728, rs61905072, rs66505542, rs7102314, rs964184, rs75198898, rs142958146, rs2075291, rs3135506, rs651821, rs45611741, rs662799, rs10750097, rs9804646, rs978880643, rs2070669, rs76353203, rs138326449, rs147210663, rs140621530, rs525028, rs141469619, rs188287950, rs202207736 |
| *CETP* | 16 | Promoter (DHS) | 411 | 1.21E-12 | 5.75E-04 | rs35571500, rs247617, rs17231506, rs34498052, rs34119551, rs34065661, rs1597000001*, rs7499892, rs5883, rs289719, rs11860407, rs189866004, rs5880 |
| *APOA1* | 11 | Enhancer (CAGE) | 642 | 1.88E-24 | 6.23E-04 | rs509728, rs61905072, rs66505542, rs7102314, rs964184, rs75198898, rs142958146, rs2075291, rs3135506, rs651821, rs45611741, rs662799, rs10750097, rs9804646, rs978880643, rs2070669, rs76353203, rs138326449, rs147210663, rs140621530, rs525028, rs141469619, rs188287950, rs202207736 |
| *SPC24* | 19 | Enhancer (CAGE) | 366 | 1.33E-08 | 4.88E-04 | rs140753491, rs138294113, rs17242353, rs17242843, rs10422256, rs72658860, rs11669576, rs2738447, rs72658867, rs2738464, rs6511728, rs3760782, rs59168178, rs2278426, rs112942459 |
| *NIPSNA P3A* | 9 | Enhancer (DHS) | 767 | 2.63E-08 | 8.46E-06 | rs2150867, rs33918808, rs112853430, rs4149307, rs9282541, rs1883025, rs1800978 |
| *LIPC* | 15 | Enhancer (DHS) | 3714 | 4.26E-08 | 1.25E-04 | rs1973688, rs1601935, rs2043082, rs10468017, rs1532085, rs436965, rs35980001, rs1800588, rs2070895, rs113298164 |
| *RP11-310H4.2* | 7 | ncRNA | 154 | 1.69E-06 | 1.69E-06 | n/a |
| *MIR4497* | 12 | ncRNA | 23 | 1.37E-06 | 1.42E-06 | rs5800864 |
| *RP11-15F12.3* | 18 | ncRNA | 64 | 7.53E-11 | 7.50E-03 | rs77960347, rs117623631, rs9958734, rs7229562, rs8086351, rs10048323, rs8084172 |

\* Samoan-specific missense variant.

MultiSTAAR-O further identified 6 genome-wide significant associations using unconditional multi-trait analysis at *α=*0.05/20,000 =2.50 ×10^−6^accounting for ncRNA genes (**Extended Data Fig. 1e, Supplementary Table 5**), with 3 rare variant associations in *RP11-15F12*.*3, RP11-310H4*.*2* and *MIR4497* remained significant at *α=*0.05/6 =8.33 ×10^−3^ after conditioning on known lipids-associated variants^26,38,39^ (**Table 2**).

Notably, among the 9 conditionally significant noncoding rare variants associations with lipid traits, 4 of them were not detected by any of the three single-trait analysis (LDL-C, HDL-C or TG) using unconditional analysis of STAAR-O, including the associations of enhancer DHS rare variants in *NIPSNAP3A* and *LIPC* as well as ncRNA rare variants in *RP11-310H4*.*2* and *MIR4497* (**Supplementary Table 5**). These results demonstrate that MultiSTAAR-O can increase power over existing methods, and identify additional trait-associated signals by leveraging cross-phenotype correlations between multiple traits.

### Genetic region multi-trait analysis of rare variants

We next applied MultiSTAAR-O to perform genetic region multi-trait analysis to identify rare variants associated with lipid traits in TOPMed. Rare variants residing in 2-kilobase (kb) sliding windows with a 1-kb skip length were aggregated and analyzed using a joint model for LDL-C, HDL-C and TG. We incorporated 12 quantitative annotations, including 9 aPCs, CADD, LINSIGHT, FATHMM-XF along with the two MAF weights in MultiSTAAR-O (**Methods**). The overall distribution of MultiSTAAR-O *P* values was well-calibrated for the multi-trait analysis (**Fig. 2b**). At a Bonferroni-corrected significance threshold of *α=*0.05/(2.65 × 10^6^) =1.89 ×10^−8^ accounting for 2.65 million 2-kb sliding windows across the genome, MultiSTAAR-O identified 502 genome-wide significant associations using unconditional multi-trait analysis (**Fig. 2a, Supplementary Table 6**). By dynamically incorporating multiple functional annotations capturing different aspects of variant function, MultiSTAAR-O detected more significant sliding windows and showed consistently smaller *P* values for top sliding windows compared with multi-trait analysis using only MAFs as the weight (**Fig. 2c**). After conditioning on known lipids-associated variants^26,38,39^, 7 out of the 502 associations remained significant at the Bonferroni-corrected threshold of *α=*0.05/502 =9.96 ×10^−5^ (**Table 3**), including two sliding windows in *DOCK7* (chromosome 1: 62,651,447 - 62,653,446 bp; chromosome 1: 62,652,447 - 62,654,446 bp) and an intergenic sliding window (chromosome 1: 145,530,447 - 145,532,446 bp) that were not detected by any of the three single-trait analysis (LDL-C, HDL-C or TG) using STAAR-O (**Supplementary Table 6**). Notably, all known lipids-associated variants indexed in the previous literature were at least 1-Mb away from the intergenic sliding window.

**Fig. 2.**
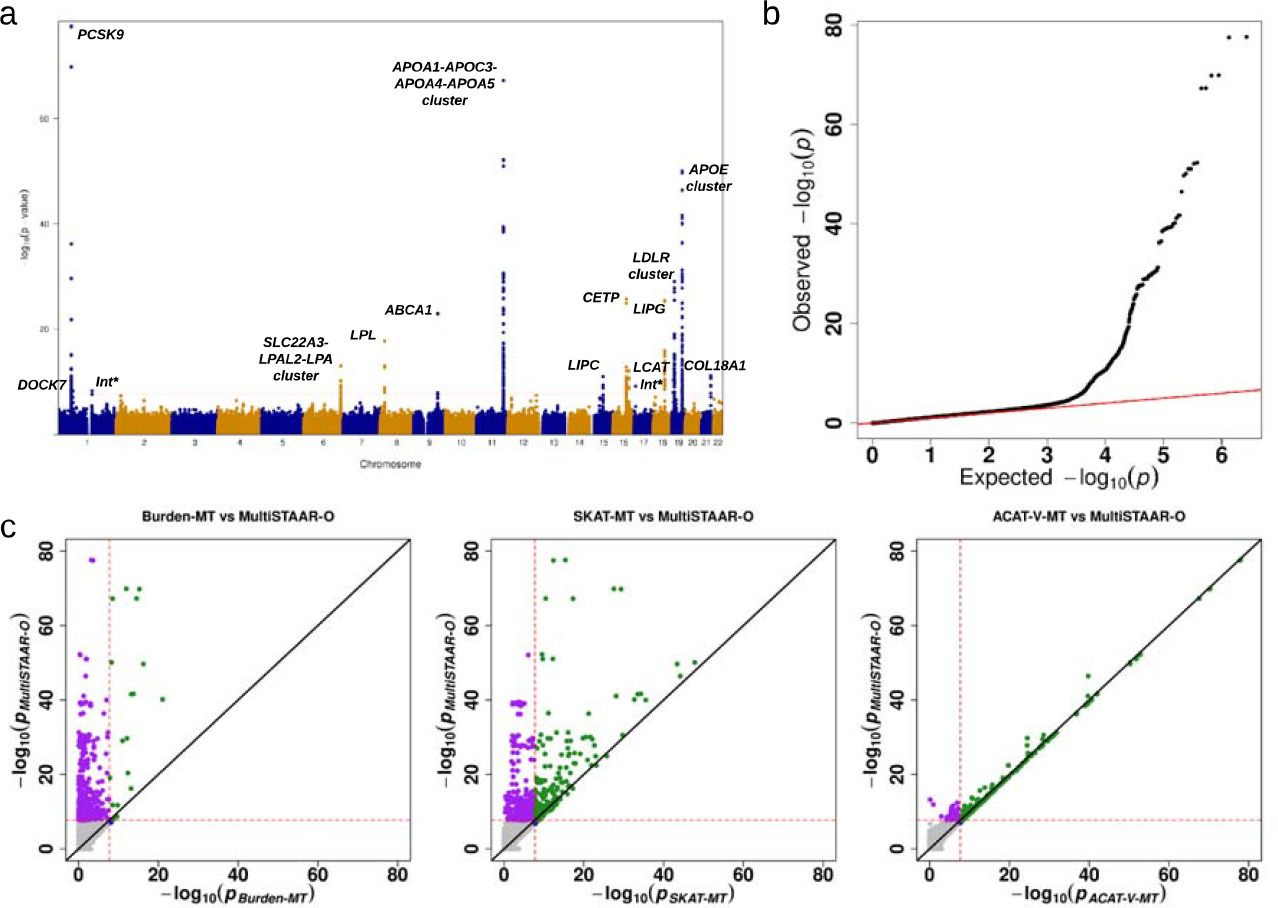
TOPMed Genetic region (2-kb sliding window) unconditional multi-trait analysis results of low-density lipoprotein cholesterol (LDL-C), high-density lipoprotein cholesterol (HDL-C) and triglycerides (TG) using TOPMed data. **a**, Manhattan plot showing the associations of 2.65 million 2-kb sliding windows versus −log_10_ (*P*) MultiSTAAR-O .The horizontal line indicates a genome-wide *P value* threshold of 1.89 ×10-8 (*n* = 61,838). **b**, Quantile-quantile plot of 2-kb sliding window MultiSTAAR-O *P* values (*n* = 61,838). **c**, Scatterplot of *P* values for the 2-kb sliding windows comparing MultiSTAAR-O with Burden-MT, SKAT-MT and ACAT-V-MT tests (MT is short for Multi-Trait). Each dot represents a sliding window with x-axis label being the −log_10_ (*P*) of the conventional multi-trait test and y-axis label being the −log_10_ (*P*) of MultiSTAAR-O (*n* = 61,838). Burden-MT, SKAT-MT, ACAT-V-MT and MultiSTAAR-O are two-sided tests. Int*, intergenic sliding window.

**Table 3.** TOPMed Genetic region (2-kb sliding window) multi-trait analysis results of both unconditional analysis and analysis conditional on known lipid-associated variants. A total of 61,838 samples from the TOPMed Program were considered in the analysis. Results for the conditionally significant sliding windows(unconditional MultiSTAAR-O *P* < 1.89 × 10^−8^and conditional MultiSTAAR-O *P* < 9.96 × 10^−5^) are presented in the table. MultiSTAAR-O is a two-sided test. Chr. no., chromosome number; Start location, start location of the 2-kb sliding window; End location, end location of the 2-kb sliding window; No. of SNVs, number of rare variants (MAF < 1%) in the 2-kb sliding window; MultiSTAAR-O, MultiSTAAR-O *P* value; Variants (adjusted), adjusted variants in the conditional analysis; n/a, no variant adjusted in the conditional analysis. Physical positions of each window are on build hg38.

| Chr. no. | Start location | End location | Gene | No. of SNVs | MultiSTAAR-O (Unconditional) | MultiSTAAR-O (Conditional) | Variants (adjusted) |
| --- | --- | --- | --- | --- | --- | --- | --- |
| 1 | 55,051,447 | 55,053,446 | *PCSK9* | 327 | 7.11E-11 | 6.60E-08 | rs12117661, rs2495491, rs11591147, rs67608943, rs72646508, rs693668, rs28362261, rs28362263, rs141502002, rs505151, rs28362286 |
| 1 | 55,052,447 | 55,054,446 | *PCSK9* | 320 | 9.37E-09 | 9.07E-06 | rs12117661, rs2495491, rs11591147, rs67608943, rs72646508, rs693668, rs28362261, rs28362263, rs141502002, rs505151, rs28362286 |
| 1 | 62,651,447 | 62,653,446 | *DOCK7* | 277 | 5.08E-09 | 7.56E-10 | rs67461605 |
| 1 | 62,652,447 | 62,654,446 | *DOCK7* | 257 | 4.87E-09 | 7.24E-10 | rs67461605 |
| 1 | 145,530,447 | 145,532,446 | *intergenic* | 233 | 5.12E-09 | 5.12E-09 | n/a |
| 19 | 11,104,367 | 11,106,366 | *LDLR* | 336 | 1.15E-12 | 8.33E-13 | rs140753491, rs138294113, rs17242353, rs17242843, rs10422256, rs72658860, rs11669576, rs2738447, rs72658867, rs2738464, rs6511728, rs3760782, rs59168178, rs2278426, rs112942459 |
| 19 | 11,105,367 | 11,107,366 | *LDLR* | 338 | 5.97E-14 | 5.55E-15 | rs140753491, rs138294113, rs17242353, rs17242843, rs10422256, rs72658860, rs11669576, rs2738447, rs72658867, rs2738464, rs6511728, rs3760782, rs59168178, rs2278426, rs112942459 |

\* Samoan-specific missense variant.

### Comparison of MultiSTAAR-O with existing multi-trait rare variant tests

Using TOPMed Freeze 8 WGS data, our gene-centric multi-trait analysis of coding rare variants identified 34 conditionally significant associations with lipid traits (**Table 1**), including *NPC1L1* and *SCARB1* missense rare variants that were missed by multi-trait burden, SKAT and ACAT-V tests (**Supplementary Table 4**). Among the 9 and 7 conditionally significant associations detected in gene-centric multi-trait analysis of noncoding rare variants and genetic region multi-trait analysis, MultiSTAAR-O identified 1 and 2 associations, respectively, that were missed by multi-trait burden, SKAT and ACAT-V tests (**Supplementary Tables 5-6**). These associations included enhancer CAGE rare variants in *SPC24* and two sliding windows in *LDLR* (chromosome 19: 11,104,367 - 11,106,366 bp; chromosome 19: 11,105,367 - 11,107,366 bp).

### Computation cost

The computational cost for MultiSTAAR-O to perform WGS multi-trait rare variant analysis of *n* = 61,838 related TOPMed lipids samples was 2 hours using 250 2.10-GHz computing cores with 12-GB memory for gene-centric coding analysis; or 20 hours using 250 2.10-GHz computing cores with 24-GB memory for gene-centric noncoding analysis; 2 hours using 250 2.10-GHz computing cores with 12-GB memory of ncRNA analysis; and 20 hours using 500 2.10-GHz computing cores with 24-GB memory for sliding window analysis. Runtime for all analyses scales linearly with the sample size^24^.

## Discussion

In this study, we have introduced MultiSTAAR as a general statistical framework and a flexible analytical pipeline for performing functionally-informed multi-trait RVAS in large-scale WGS studies. MultiSTAAR improves power by analyzing multiple traits simultaneously and dynamically incorporating multiple functional annotations, while accounting for relatedness and population structure among study samples.

By jointly analyzing multiple quantitative traits using a multivariate linear mixed model, MultiSTAAR explicitly leverages the correlation among multiple phenotypes to enhance power for detecting additional association signals, outperforming single-trait analyses of the individual phenotypes. MultiSTAAR also enables conditional multi-trait analysis to identify putatively novel rare variant associations independent of a set of known variants. Using TOPMed Freeze 8 WGS data, our gene-centric multi-trait analysis of noncoding rare variants identified 9 conditionally significant associations with lipid traits (**Table 2**), including 4 noncoding associations that were missed by single-trait analysis using STAAR (**Supplementary Table 5**). Our genetic region multi-trait analysis of rare variants identified 7 conditionally significant 2-kb sliding windows associated with lipid traits (**Table 3**), including 3 associations that were missed by single-trait analysis using STAAR (**Supplementary Table 6**).

By dynamically incorporating multiple annotations capturing diverse aspects of variant biological function in the second step, MultiSTAAR further improves power over existing multi-trait rare variant analysis methods. Our simulation studies demonstrated that MultiSTAAR-O maintained accurate type I error rates while achieving considerable power gains over multi-trait burden, SKAT and ACAT-V tests that do not incorporate functional annotation information (**Extended Data Figs. 2-5, Supplementary Figs. 1-4**). Notably, the existing ACAT-V method^9^ does not support multi-trait analysis. We extended it to accommodate multi-trait settings and incorporated the multi-trait ACAT-V test into the MultiSTAAR framework (**Methods**).

Implemented as a flexible analytical pipeline, MultiSTAAR allows for customized input phenotype selection, variant set definition and user-specified annotation weights to facilitate functionally-informed multi-trait analyses. In addition to rare variant association analysis of coding and noncoding regions, MultiSTAAR also provides single-variant multi-trait analysis for common and low-frequency variants under a given MAF or minor allele count (MAC) cutoff (e.g. MAC ≥ 20). Using 61,838 TOPMed lipids samples, it took 8 hours using 250 2.10-GHz computing cores with 12-GB memory for single-variant multi-trait analysis, which is scalable for large WGS/WES datasets. On the other hand, MultiSTAAR could be further extended to allow for dynamic windows with data-adaptive sizes in genetic region analysis^24,44^, to properly leverage synthetic surrogates in the presence of partially missing phenotypes^45^, and to incorporate summary statistics for meta-analysis of multiple WGS/WES studies^46^.

In summary, MultiSTAAR provides a powerful statistical framework and a computationally scalable analytical pipeline for large-scale WGS multi-trait analysis with complex study samples. Compared to single-trait analysis, MultiSTAAR offers a notable increase in statistical power when analyzing multiple moderately to highly correlated traits, all while maintaining control over type I error rates across various genetic architectures. As the sample sizes and number of available phenotypes increase in biobank-scale sequencing studies, our proposed method may contribute to a better understanding of the genetic architecture of complex traits by elucidating the role of rare variants with pleiotropic effects.

## Acknowledgments

This work was supported by grants R35-CA197449, U19-CA203654, U01-HG012064, and U01-HG009088 (X. Lin), NHLBI TOPMed Fellowship (X. Li), R01-HL142711 and R01-HL127564 (P.N. and G.M.P.), 75N92020D00001, HHSN268201500003I, N01-HC-95159, 75N92020D00005, N01-HC-95160, 75N92020D00002, N01-HC-95161, 75N92020D00003, N01-HC-95162, 75N92020D00006, N01-HC-95163, 75N92020D00004, N01-HC-95164, 75N92020D00007, N01-HC-95165, N01-HC-95166, N01-HC-95167, N01-HC-95168, N01-HC-95169, UL1-TR-000040, UL1-TR-001079, UL1-TR-001420, UL1-TR001881, DK063491, R01-HL071051, R01-HL071205, R01-HL071250, R01-HL071251, R01-HL071258, R01-HL071259, and UL1-RR033176 (J.I.R.), HHSN268201800001I and U01-HL137162 (K.M.R.), 1R35-HL135818, R01-HL113338, and HL046389 (S.R.), HL105756 (B.M.P.), HHSN268201600018C, HHSN268201600001C, HHSN268201600002C, HHSN268201600003C, and HHSN268201600004C (C.K.), R01-MD012765 and R01-DK117445 (N.F.), 18CDA34110116 from American Heart Association (P.S.d.V.), R01-HL153805, R03-HL154284 (B.E.C.), HHSN268201700001I, HHSN268201700002I, HHSN268201700003I, HHSN268201700005I, and HHSN268201700004I (E.B.), U01-HL072524, R01-HL104135-04S1, U01-HL054472, U01-HL054473, U01-HL054495, U01-HL054509, and R01-HL055673-18S1 (D.K.A.), U01-HL72518, HL087698, HL49762, HL59684, HL58625, HL071025, HL112064, NR0224103, and M01-RR000052 (to the Johns Hopkins General Clinical Research Center). This work was supported by R01 HL92301, R01 HL67348, R01 NS058700, R01 AR48797, R01 DK071891, R01 AG058921, the General Clinical Research Center of the Wake Forest University School of Medicine (M01 RR07122, F32 HL085989), the American Diabetes Association, and a pilot grant from the Claude Pepper Older Americans Independence Center of Wake Forest University Health Sciences (P60 AG10484). The Framingham Heart Study (FHS) acknowledges the support of contracts NO1-HC-25195, HHSN268201500001I and 75N92019D00031 from the National Heart, Lung and Blood Institute and grant supplement R01 HL092577-06S1 for this research. We also acknowledge the dedication of the FHS study participants without whom this research would not be possible. R.S.V. is supported in part by the Evans Medical Foundation and the Jay and Louis Coffman Endowment from the Department of Medicine, Boston University School of Medicine. The Jackson Heart Study (JHS) is supported and conducted in collaboration with Jackson State University (HHSN268201800013I), Tougaloo College (HHSN268201800014I), the Mississippi State Department of Health (HHSN268201800015I) and the University of Mississippi Medical Center (HHSN268201800010I, HHSN268201800011I and HHSN268201800012I) contracts from the National Heart, Lung, and Blood Institute (NHLBI) and the National Institute on Minority Health and Health Disparities (NIMHD). The authors also wish to thank the staffs and participants of the JHS. Support for GENOA was provided by the National Heart, Lung and Blood Institute (U01HL054457, U01HL054464, U01HL054481, R01HL119443, and R01HL087660) of the National Institutes of Health. Collection of the San Antonio Family Study data was supported in part by National Institutes of Health (NIH) grants P01 HL045522, MH078143, MH078111 and MH083824; and whole genome sequencing of SAFS subjects was supported by U01 DK085524 and R01 HL113323. Molecular data for the Trans-Omics in Precision Medicine (TOPMed) program was supported by the National Heart, Lung and Blood Institute (NHLBI). Core support including centralized genomic read mapping and genotype calling, along with variant quality metrics and filtering were provided by the TOPMed Informatics Research Center (3R01HL-117626-02S1; contract HHSN268201800002I). Core support including phenotype harmonization, data management, sample-identity QC, and general program coordination were provided by the TOPMed Data Coordinating Center (R01HL-120393; U01HL-120393; contract HHSN268201800001I). We gratefully acknowledge the studies and participants who provided biological samples and data for TOPMed. The full study specific acknowledgements are detailed in **Supplementary Note**.

## Author contributions

X. Li, H.C., Z. Li, Z. Liu and X. Lin designed the experiments. X. Li, H.C., Z. Li and X. Lin performed the experiments. X. Li, H.C., M.S.S., E.V.B., Y.W., R.S., Z.R.M., Z.Y., D.K.A., J.C.B., J.B., E.B., D.W.B., J.A.B., B.E.C., A.P.C., J.C.C., N.C., Y.D.I.C., J.E.C., P.S.d.V., M.F., N.F., B.I.F., C.G., N.L.H.C., J.H., L.H., Y.J.H., M.R.I., R.C.K., S.L.R.K., T.K., I.K., C.K., B.G.K., C.L., R.J.F.L., M.C.M., L.W.M., R.A.M., R.L.M., B.D.M., M.E.M., A.C.M., N.D.P., P.A.P., B.M.P., L.M.R., S.R., A.P.R., S.S.R., C.M.S., J.A.S., K.D.T., H.T., R.S.V., Z.W., L.R.Y., B.Y., K.M.R., J.I.R., G.M.P., P.N., Z. Li, Z. Liu and X. Lin acquired, analyzed or interpreted data. G.M.P., P.N. and the NHLBI TOPMed Lipids Working Group provided administrative, technical or material support. X. Li, Z. Li, Z. Liu and X. Lin drafted the manuscript and revised it according to suggestions by the coauthors. All authors critically reviewed the manuscript, suggested revisions as needed and approved the final version.

## Competing interests

Z.R.M. is an employee of Insitro. M.E.M. receives research funding from Regeneron Pharmaceutical Inc., unrelated to this project. B.M.P. serves on the Steering Committee of the Yale Open Data Access Project funded by Johnson & Johnson. L.M.R. is a consultant for the TOPMed Administrative Coordinating Center (via Westat). X. Lin is a consultant of AbbVie Pharmaceuticals and Verily Life Sciences. The remaining authors declare no competing interests.

### NHLBI Trans-Omics for Precisition Medicine (TOPMed) Consortium

Namiko Abe^50^, Gonçalo Abecasis^51^, Francois Aguet^52^, Christine Albert^53^, Laura Almasy^54^, Alvaro Alonso^55^, Seth Ament^56^, Peter Anderson^57^, Pramod Anugu^58^, Deborah Applebaum-Bowden^59^, Kristin Ardlie^52^, Dan Arking^60^, Allison Ashley-Koch^61^, Stella Aslibekyan^62^, Tim Assimes^63^, Paul Auer^64^, Dimitrios Avramopoulos^60^, Najib Ayas^65^, Adithya Balasubramanian^66^, John Barnard^67^, Kathleen Barnes^68^, R. Graham Barr^69^, Emily Barron-Casella^60^, Lucas Barwick^70^, Terri Beaty^60^, Gerald Beck^71^, Diane Becker^72^, Lewis Becker^60^, Rebecca Beer^73^, Amber Beitelshees^56^, Emelia Benjamin^74^, Takis Benos^75^, Marcos Bezerra^76^, Larry Bielak^51^, Thomas Blackwell^51^, Nathan Blue^77^, Russell Bowler^78^, Ulrich Broeckel^79^, Jai Broome^57^, Deborah Brown^80^, Karen Bunting^50^, Esteban Burchard^81^, Carlos Bustamante^82^, Erin Buth^83^, Jonathan Cardwell^84^, Vincent Carey^85^, Julie Carrier^86^, Cara Carty^87^, Richard Casaburi^88^, Juan P Casas Romero^89^, James Casella^60^, Peter Castaldi^90^, Mark Chaffin^52^, Christy Chang^56^, Yi-Cheng Chang^91^, Daniel Chasman^92^, Sameer Chavan^84^, Bo-Juen Chen^50^, Wei-Min Chen^93^, Michael Cho^85^, Seung Hoan Choi^52^, Lee-Ming Chuang^94^, Mina Chung^95^, Ren-Hua Chung^96^, Clary Clish^97^, Suzy Comhair^98^, Matthew Conomos^83^, Elaine Cornell^99^, Adolfo Correa^100^, Carolyn Crandall^88^, James Crapo^101^, L. Adrienne Cupples^102^, Jeffrey Curtis^103^, Brian Custer^104^, Coleen Damcott^56^, Dawood Darbar^105^, Sean David^106^, Colleen Davis^57^, Michelle Daya^84^, Michael DeBaun^107^, Dawn DeMeo^85^, Ranjan Deka^108^, Scott Devine^56^, Huyen Dinh^66^, Harsha Doddapaneni^109^, Qing Duan^110^, Shannon Dugan-Perez^111^, Ravi Duggirala^112^, Jon Peter Durda^113^, Susan K. Dutcher^114^, Charles Eaton^115^, Lynette Ekunwe^58^, Adel El Boueiz^116^, Patrick Ellinor^117^, Leslie Emery^57^, Serpil Erzurum^118^, Charles Farber^93^, Jesse Farek^66^, Tasha Fingerlin^119^, Matthew Flickinger^51^, Chris Frazar^57^, Mao Fu^56^, Stephanie M. Fullerton^57^, Lucinda Fulton^120^, Stacey Gabriel^52^, Weiniu Gan^73^, Shanshan Gao^84^, Yan Gao^58^, Margery Gass^121^, Heather Geiger^122^, Bruce Gelb^123^, Mark Geraci^124^, Soren Germer^50^, Robert Gerszten^125^, Auyon Ghosh^85^, Richard Gibbs^66^, Chris Gignoux^63^, Mark Gladwin^126^, David Glahn^127^, Stephanie Gogarten^57^, Da-Wei Gong^56^, Harald Goring^128^, Sharon Graw^129^, Kathryn J. Gray^130^, Daniel Grine^84^, Colin Gross^51^, Yue Guan^56^, Xiuqing Guo^131^, Namrata Gupta^132^, Jeff Haessler^121^, Michael Hall^133^, Yi Han^66^, Patrick Hanly^134^, Daniel Harris^135^, Nicola L. Hawley^136^, Ben Heavner^83^, Susan Heckbert^137^, Ryan Hernandez^81^, David Herrington^138^, Craig Hersh^139^, Bertha Hidalgo^62^, James Hixson^140^, Brian Hobbs^141^, John Hokanson^84^, Elliott Hong^56^, Karin Hoth^142^, Chao (Agnes) Hsiung^143^, Jianhong Hu^66^, Haley Huston^144^, Chii Min Hwu^145^, Rebecca Jackson^146^, Deepti Jain^57^, Cashell Jaquish^147^, Jill Johnsen^148^, Andrew Johnson^73^, Craig Johnson^57^, Rich Johnston^55^, Kimberly Jones^60^, Hyun Min Kang^149^, Shannon Kelly^150^, Eimear Kenny^123^, Michael Kessler^56^, Alyna Khan^57^, Ziad Khan^66^, Wonji Kim^151^, John Kimoff^152^, Greg Kinney^153^, Barbara Konkle^154^, Holly Kramer^155^, Christoph Lange^156^, Ethan Lange^84^, Leslie Lange^157^, Cathy Laurie^57^, Cecelia Laurie^57^, Meryl LeBoff^85^, Jonathon LeFaive^51^, Jiwon Lee^85^, Sandra Lee^66^, Wen-Jane Lee^145^, David Levine^57^, Daniel Levy^73^, Joshua Lewis^56^, Xiaohui Li^131^, Yun Li^110^, Henry Lin^131^, Honghuang Lin^158^, Simin Liu^159^, Yongmei Liu^160^, Yu Liu^161^, Steven Lubitz^117^, Kathryn Lunetta^162^, James Luo^73^, Ulysses Magalang^163^, Barry Make^60^, Ani Manichaikul^93^, Alisa Manning^164^, JoAnn Manson^85^, Melissa Marton^122^, Susan Mathai^84^, Susanne May^83^, Patrick McArdle^56^, Merry-Lynn McDonald^165^, Sean McFarland^151^, Stephen McGarvey^166^, Daniel McGoldrick^167^, Caitlin McHugh^83^, Becky McNeil^168^, Hao Mei^58^, James Meigs^169^, Vipin Menon^66^, Luisa Mestroni^129^, Ginger Metcalf^66^, Deborah A Meyers^170^, Emmanuel Mignot^171^, Julie Mikulla^73^, Nancy Min^58^, Mollie Minear^172^, Matt Moll^90^, Zeineen Momin^66^, Courtney Montgomery^173^, Donna Muzny^66^, Josyf C Mychaleckyj^93^, Girish Nadkarni^123^, Rakhi Naik^60^, Take Naseri^174^, Sergei Nekhai^175^, Sarah C. Nelson^83^, Bonnie Neltner^84^, Caitlin Nessner^66^, Deborah Nickerson^176^, Osuji Nkechinyere^66^, Kari North^110^, Jeff O’Connell^177^, Tim O’Connor^56^, Heather Ochs-Balcom^178^, Geoffrey Okwuonu^66^, Allan Pack^179^, David T. Paik^180^, James Pankow^181^, George Papanicolaou^73^, Cora Parker^182^, Juan Manuel Peralta^112^, Marco Perez^63^, James Perry^56^, Ulrike Peters^183^, Lawrence S Phillips^55^, Jacob Pleiness^51^, Toni Pollin^56^, Wendy Post^184^, Julia Powers Becker^185^, Meher Preethi Boorgula^84^, Michael Preuss^123^, Pankaj Qasba^73^, Dandi Qiao^85^, Zhaohui Qin^55^, Nicholas Rafaels^186^, Mahitha Rajendran^66^, D.C. Rao^120^, Laura Rasmussen-Torvik^187^, Aakrosh Ratan^93^, Robert Reed^56^, Catherine Reeves^188^, Elizabeth Regan^189^, Muagututi’a Sefuiva Reupena^190^, Rebecca Robillard^191^, Nicolas Robine^122^, Dan Roden^192^, Carolina Roselli^52^, Ingo Ruczinski^60^, Alexi Runnels^122^, Pamela Russell^84^, Sarah Ruuska^193^, Kathleen Ryan^56^, Ester Cerdeira Sabino^194^, Danish Saleheen^195^, Shabnam Salimi^196^, Sejal Salvi^66^, Steven Salzberg^60^, Kevin Sandow^197^, Vijay G. Sankaran^198^, Jireh Santibanez^66^, Karen Schwander^120^, David Schwartz^84^, Frank Sciurba^126^, Christine Seidman^199^, Jonathan Seidman^200^, Vivien Sheehan^201^, Stephanie L. Sherman^202^, Amol Shetty^56^, Aniket Shetty^84^, Wayne Hui-Heng Sheu^145^, M. Benjamin Shoemaker^203^, Brian Silver^204^, Edwin Silverman^85^, Robert Skomro^205^, Albert Vernon Smith^206^, Josh Smith^57^, Nicholas Smith^207^, Tanja Smith^50^, Sylvia Smoller^208^, Beverly Snively^209^, Michael Snyder^210^, Tamar Sofer^125^, Nona Sotoodehnia^57^, Adrienne M. Stilp^57^, Garrett Storm^211^, Elizabeth Streeten^56^, Jessica Lasky Su^212^, Yun Ju Sung^120^, Jody Sylvia^85^, Adam Szpiro^57^, Frédéric Sériès^213^, Daniel Taliun^51^, Hua Tang^210^, Margaret Taub^60^, Matthew Taylor^129^, Simeon Taylor^56^, Marilyn Telen^61^, Timothy A. Thornton^57^, Machiko Threlkeld^214^, Lesley Tinker^215^, David Tirschwell^57^, Sarah Tishkoff^216^, Catherine Tong^217^, Russell Tracy^218^, Michael Tsai^181^, Dhananjay Vaidya^60^, David Van Den Berg^219^, Peter VandeHaar^51^, Scott Vrieze^181^, Tarik Walker^84^, Robert Wallace^142^, Avram Walts^84^, Fei Fei Wang^57^, Heming Wang^220^, Jiongming Wang^221^, Karol Watson^88^, Jennifer Watt^66^, Daniel E. Weeks^222^, Joshua Weinstock^149^, Bruce Weir^57^, Scott T Weiss^223^, Lu-Chen Weng^117^, Jennifer Wessel^224^, Cristen Willer^103^, Kayleen Williams^83^, L. Keoki Williams^225^, Scott Williams^226^, Carla Wilson^85^, James Wilson^227^, Lara Winterkorn^122^, Quenna Wong^57^, Baojun Wu^228^, Joseph Wu^180^, Huichun Xu^56^, Ivana Yang^84^, Ketian Yu^51^, Seyedeh Maryam Zekavat^52^, Yingze Zhang^229^, Snow Xueyan Zhao^101^, Wei Zhao^230^, Xiaofeng Zhu^231^, Elad Ziv^232^, Michael Zody^50^, Sebastian Zoellner^51^, Mariza de Andrade^233^, Lisa de las Fuentes^234^

50 - New York Genome Center, New York, New York, 10013, US; 51 - University of Michigan, Ann Arbor, Michigan, 48109, US; 52 - Broad Institute, Cambridge, Massachusetts, 2142, US; 53 - Cedars Sinai, Boston, Massachusetts, 2114, US; 54 - Children’s Hospital of Philadelphia, University of Pennsylvania, Philadelphia, Pennsylvania, 19104, US; 55 - Emory University, Atlanta, Georgia, 30322, US; 56 - University of Maryland, Baltimore, Maryland, 21201, US; 57 - University of Washington, Seattle, Washington, 98195, US; 58 - University of Mississippi, Jackson, Mississippi, 38677, US; 59 - National Institutes of Health, Bethesda, Maryland, 20892, US; 60 - Johns Hopkins University, Baltimore, Maryland, 21218, US; 61 - Duke University, Durham, North Carolina, 27708, US; 62 - University of Alabama, Birmingham, Alabama, 35487, US; 63 - Stanford University, Stanford, California, 94305, US; 64 - Medical College of Wisconsin, Milwaukee, Wisconsin, 53211, US; 65 - Providence Health Care, Medicine, Vancouver, CA; 66 - Baylor College of Medicine Human Genome Sequencing Center, Houston, Texas, 77030, US; 67 - Cleveland Clinic, Cleveland, Ohio, 44195, US; 68 - Tempus, University of Colorado Anschutz Medical Campus, Aurora, Colorado, 80045, US; 69 - Columbia University, New York, New York, 10032, US; 70 - The Emmes Corporation, LTRC, Rockville, Maryland, 20850, US; 71 - Cleveland Clinic, Quantitative Health Sciences, Cleveland, Ohio, 44195, US; 72 - Johns Hopkins University, Medicine, Baltimore, Maryland, 21218, US; 73 - National Heart, Lung, and Blood Institute, National Institutes of Health, Bethesda, Maryland, 20892, US; 74 - Boston University, Massachusetts General Hospital, Boston University School of Medicine, Boston, Massachusetts, 2114, US; 75 - University of Florida, Epidemiology, Gainesville, Florida, 32610, US; 76 - Fundação de Hematologia e Hemoterapia de Pernambuco - Hemope, Recife, 52011-000, BR; 77 - University of Utah, Obstetrics and Gynecology, Salt Lake City, Utah, 84132, US; 78 - National Jewish Health, National Jewish Health, Denver, Colorado, 80206, US; 79 - Medical College of Wisconsin, Pediatrics, Milwaukee, Wisconsin, 53226, US; 80 - University of Texas Health at Houston, Pediatrics, Houston, Texas, 77030, US; 81 - University of California, San Francisco, San Francisco, California, 94143, US; 82 - Stanford University, Biomedical Data Science, Stanford, California, 94305, US; 83 - University of Washington, Biostatistics, Seattle, Washington, 98195, US; 84 - University of Colorado at Denver, Denver, Colorado, 80204, US; 85 - Brigham & Women’s Hospital, Boston, Massachusetts, 2115, US; 86 - University of Montreal, US; 87 - Washington State University, Pullman, Washington, 99164, US; 88 - University of California, Los Angeles, Los Angeles, California, 90095, US; 89 - Brigham & Women’s Hospital, US; 90 - Brigham & Women’s Hospital, Medicine, Boston, Massachusetts, 2115, US; 91 - National Taiwan University, Taipei, 10617, TW; 92 - Brigham & Women’s Hospital, Division of Preventive Medicine, Boston, Massachusetts, 2215, US; 93 - University of Virginia, Charlottesville, Virginia, 22903, US; 94 - National Taiwan University, National Taiwan University Hospital, Taipei, 10617, TW; 95 - Cleveland Clinic, Cleveland Clinic, Cleveland, Ohio, 44195, US; 96 - National Health Research Institute Taiwan, Miaoli County, 350, TW; 97 - Broad Institute, Metabolomics Platform, Cambridge, Massachusetts, 2142, US; 98 - Cleveland Clinic, Immunity and Immunology, Cleveland, Ohio, 44195, US; 99 - University of Vermont, Burlington, Vermont, 5405, US; 100 - University of Mississippi, Population Health Science, Jackson, Mississippi, 39216, US; 101 - National Jewish Health, Denver, Colorado, 80206, US; 102 - Boston University, Biostatistics, Boston, Massachusetts, 2115, US; 103 - University of Michigan, Internal Medicine, Ann Arbor, Michigan, 48109, US; 104 - Vitalant Research Institute, San Francisco, California, 94118, US; 105 - University of Illinois at Chicago, Chicago, Illinois, 60607, US; 106 - University of Chicago, Chicago, Illinois, 60637, US; 107 - Vanderbilt University, Nashville, Tennessee, 37235, US; 108 - University of Cincinnati, Cincinnati, Ohio, 45220, US; 109 - Baylor College of Medicine Human Genome Sequencing Center, Houston, Texas, 77030; 110 - University of North Carolina, Chapel Hill, North Carolina, 27599, US; 111 - Baylor College of Medicine Human Genome Sequencing Center, BCM, Houston, Texas, 77030, US; 112 - University of Texas Rio Grande Valley School of Medicine, Edinburg, Texas, 78539, US; 113 - University of Vermont, Pathology and Laboratory Medicine, Burlington, Vermont, 5405, US; 114 - Washington University in St Louis, Genetics, St Louis, Missouri, 63110, US; 115 - Brown University, Providence, Rhode Island, 2912, US; 116 - Harvard University, Channing Division of Network Medicine, Cambridge, Massachusetts, 2138, US; 117 - Massachusetts General Hospital, Boston, Massachusetts, 2114, US; 118 - Cleveland Clinic, Lerner Research Institute, Cleveland, Ohio, 44195, US; 119 - National Jewish Health, Center for Genes, Environment and Health, Denver, Colorado, 80206, US; 120 - Washington University in St Louis, St Louis, Missouri, 63130, US; 121 - Fred Hutchinson Cancer Research Center, Seattle, Washington, 98109, US; 122 - New York Genome Center, New York City, New York, 10013, US; 123 - Icahn School of Medicine at Mount Sinai, New York, New York, 10029, US; 124 - University of Pittsburgh, Pittsburgh, Pennsylvania, US; 125 Beth Israel Deaconess Medical Center, Boston, Massachusetts, 2215, US; 126 - University of Pittsburgh, Pittsburgh, Pennsylvania, 15260, US; 127 - Boston Children’s Hospital, Harvard Medical School, Department of Psychiatry, Boston, Massachusetts, 2115, US; 128 - University of Texas Rio Grande Valley School of Medicine, San Antonio, Texas, 78229, US; 129 - University of Colorado Anschutz Medical Campus, Aurora, Colorado, 80045, US; 130 - Mass General Brigham, Obstetrics and Gynecology, Boston, Massachusetts, 2115, US; 131 - Lundquist Institute, Torrance, California, 90502, US; 132 - Broad Institute, Broad Institute, Cambridge, Massachusetts, 2142, US; 133 - University of Mississippi, Cardiology, Jackson, Mississippi, 39216, US; 134 - University of Calgary, Medicine, Calgary, CA; 135 - University of Maryland, Genetics, Philadelphia, Pennsylvania, 19104, US; 136 - Yale University, Department of Chronic Disease Epidemiology, New Haven, Connecticut, 6520, US; 137 - University of Washington, Epidemiology, Seattle, Washington, 98195- 9458, US; 138 - Wake Forest Baptist Health, Winston-Salem, North Carolina, 27157, US; 139 - Brigham & Women’s Hospital, Channing Division of Network Medicine, Boston, Massachusetts, 2115, US; 140 - University of Texas Health at Houston, Houston, Texas, 77225, US; 141 - Regeneron Genetics Center, Boston, Massachusetts, 2115, US; 142 - University of Iowa, Iowa City, Iowa, 52242, US; 143 - National Health Research Institute Taiwan, Institute of Population Health Sciences, NHRI, Miaoli County, 350, TW; 144 - Blood Works Northwest, Seattle, Washington, 98104, US; 145 - Taichung Veterans General Hospital Taiwan, Taichung City, 407, TW; 146 - Oklahoma State University Medical Center, Internal Medicine, DIvision of Endocrinology, Diabetes and Metabolism, Columbus, Ohio, 43210, US; 147 - National Heart, Lung, and Blood Institute, National Institutes of Health, NHLBI, Bethesda, Maryland, 20892, US; 148 - University of Washington, Medicine, Seattle, Washington, 98109, US; 149 - University of Michigan, Biostatistics, Ann Arbor, Michigan, 48109, US; 150 - University of California, San Francisco, San Francisco, California, 94118, US; 151 Harvard University, Cambridge, Massachusetts, 2138, US; 152 - McGill University, Montréal, QC H3A 0G4, CA; 153 - University of Colorado at Denver, Epidemiology, Aurora, Colorado, 80045, US; 154 - Blood Works Northwest, Medicine, Seattle, Washington, 98104, US; 155 - Loyola University, Public Health Sciences, Maywood, Illinois, 60153, US; 156 - Harvard School of Public Health, Biostats, Boston, Massachusetts, 2115, US; 157 - University of Colorado at Denver, Medicine, Aurora, Colorado, 80048, US; 158 - Boston University, University of Massachusetts Chan Medical School, Worcester, Massachusetts, 1655, US; 159 - Brown University, Epidemiology and Medicine, Providence, Rhode Island, 2912, US; 160 - Duke University, Cardiology, Durham, North Carolina, 27708, US; 161 - Stanford University, Cardiovascular Institute, Stanford, California, 94305, US; 162 - Boston University, Boston, Massachusetts, 2215, US; 163 - The Ohio State University, Division of Pulmonary, Critical Care and Sleep Medicine, Columbus, Ohio, 43210, US; 164 - Broad Institute, Harvard University, Massachusetts General Hospital; 165 - University of Alabama, University of Alabama at Birmingham, Birmingham, Alabama, 35487, US; 166 -Brown University, Epidemiology, Providence, Rhode Island, 2912, US; 167 – University of Washington, Genome Sciences, Seattle, Washington, 98195, US; 168 - RTI International, US; 169 - Massachusetts General Hospital, Medicine, Boston, Massachusetts, 2114, US; 170 - University of Arizona, Tucson, Arizona, 85721, US; 171 - Stanford University, Center For Sleep Sciences and Medicine, Palo Alto, California, 94304, US; 172 - National Institute of Child Health and Human Development, National Institutes of Health, Bethesda, Maryland, 20892, US; 173 - Oklahoma Medical Research Foundation, Genes and Human Disease, Oklahoma City, Oklahoma, 73104, US; 174 - Ministry of Health, Government of Samoa, Apia, WS; 175 - Howard University, Washington, District of Columbia, 20059, US; 176 - University of Washington, Department of Genome Sciences, Seattle, Washington, 98195, US; 177 - University of Maryland, Balitmore, Maryland, 21201, US; 178 - University at Buffalo, Buffalo, New York, 14260, US; 179 - University of Pennsylvania, Division of Sleep Medicine/Department of Medicine, Philadelphia, Pennsylvania, 19104-3403, US; 180 - Stanford University, Stanford Cardiovascular Institute, Stanford, California, 94305, US; 181 - University of Minnesota, Minneapolis, Minnesota, 55455, US; 182 - RTI International, Biostatistics and Epidemiology Division, Research Triangle Park, North Carolina, 27709-2194, US; 183 - Fred Hutchinson Cancer Research Center, Fred Hutch and UW, Seattle, Washington, 98109, US; 184 - Johns Hopkins University, Cardiology/Medicine, Baltimore, Maryland, 21218, US; 185 - University of Colorado at Denver, Medicine, Denver, Colorado, 80204, US; 186 - University of Colorado at Denver, CCPM, Denver, Colorado, 80045, US; 187 - Northwestern University, Chicago, Illinois, 60208, US; 188 - New York Genome Center, New York Genome Center, New York City, New York, 10013, US; 189 - National Jewish Health, Medicine, Denver, Colorado, 80206, US; 190 - Lutia I Puava Ae Mapu I Fagalele, Apia, WS; 191 - University of Ottawa, Sleep Research Unit, University of Ottawa Institute for Mental Health Research, Ottawa, ON K1Z 7K4, CA; 192 - Vanderbilt University, Medicine, Pharmacology, Biomedicla Informatics, Nashville, Tennessee, 37235, US; 193 - University of Washington, Seattle, Washington, 98104, US; 194 - Universidade de Sao Paulo, Faculdade de Medicina, Sao Paulo, 1310000, BR; 195 - Columbia University, New York, New York, 10027, US; 196 - University of Maryland, Pathology, Seattle, Washington, 98195, US; 197 - Lundquist Institute, TGPS, Torrance, California, 90502, US; 198 - Harvard University, Division of Hematology/Oncology, Boston, Massachusetts, 2115, US; 199 - Harvard Medical School, Genetics, Boston, Massachusetts, 2115, US; 200 - Harvard Medical School, Boston, Massachusetts, 2115, US; 201 - Emory University, Pediatrics, Atlanta, Georgia, 30307, US; 202 - Emory University, Human Genetics, Atlanta, Georgia, 30322, US; 203 - Vanderbilt University, Medicine/Cardiology, Nashville, Tennessee, 37235, US; 204 - UMass Memorial Medical Center, Worcester, Massachusetts, 1655, US; 205 - University of Saskatchewan, Saskatoon, SK S7N 5C9, CA; 206 - University of Michigan; 207 - University of Washington, Epidemiology, Seattle, Washington, 98195, US; 208 - Albert Einstein College of Medicine, New York, New York, 10461, US; 209 - Wake Forest Baptist Health, Biostatistical Sciences, Winston-Salem, North Carolina, 27157, US; 210 - Stanford University, Genetics, Stanford, California, 94305, US; 211 - University of Colorado at Denver, Genomic Cardiology, Aurora, Colorado, 80045, US; 212 - Brigham & Women’s Hospital, Channing Department of Medicine, Boston, Massachusetts, 2115, US; 213 - Université Laval, Quebec City, G1V 0A6, CA; 214 - University of Washington, University of Washington, Department of Genome Sciences, Seattle, Washington, 98195, US; 215 - Fred Hutchinson Cancer Research Center, Cancer Prevention Division of Public Health Sciences, Seattle, Washington, 98109, US; 216 - University of Pennsylvania, Genetics, Philadelphia, Pennsylvania, 19104, US; 217 - University of Washington, Department of Biostatistics, Seattle, Washington, 98195, US; 218 - University of Vermont, Pathology & Laboratory Medicine, Burlington, Vermont, 5405, US; 219 - University of Southern California, USC Methylation Characterization Center, University of Southern California, California, 90033, US; 220 - Brigham & Women’s Hospital, Mass General Brigham, Boston, Massachusetts, 2115, US; 221 - University of Michigan, US; 222 - University of Pittsburgh, Department of Human Genetics, Pittsburgh, Pennsylvania, 15260, US; 223 - Brigham & Women’s Hospital, Channing Division of Network Medicine, Department of Medicine, Boston, Massachusetts, 2115, US; 224 - Indiana University, Epidemiology, Indianapolis, Indiana, 46202, US; 225 - Henry Ford Health System, Detroit, Michigan, 48202, US; 226 - Case Western Reserve University; 227 - Beth Israel Deaconess Medical Center, Cardiology, Cambridge, Massachusetts, 2139, US; 228 - Henry Ford Health System, Department of Medicine, Detroit, Michigan, 48202, US; 229 - University of Pittsburgh, Medicine, Pittsburgh, Pennsylvania, 15260, US; 230 - University of Michigan, Department of Epidemiology, Ann Arbor, Michigan, 48109, US; 231 - Case Western Reserve University, Department of Population and Quantitative Health Sciences, Cleveland, Ohio, 44106, US; 232 - University of California, San Francisco, Medicine, San Francisco, California, 94143, US; 233 - Mayo Clinic, Health Quantitative Sciences Research, Rochester, Minnesota, 55905, US; 234 - Washington University in St Louis, Department of Medicine, Cardiovascular Division, St. Louis, Missouri, 63110, US

## EXTENDED DATA FIGURES

**Extended Data Fig. 1.**
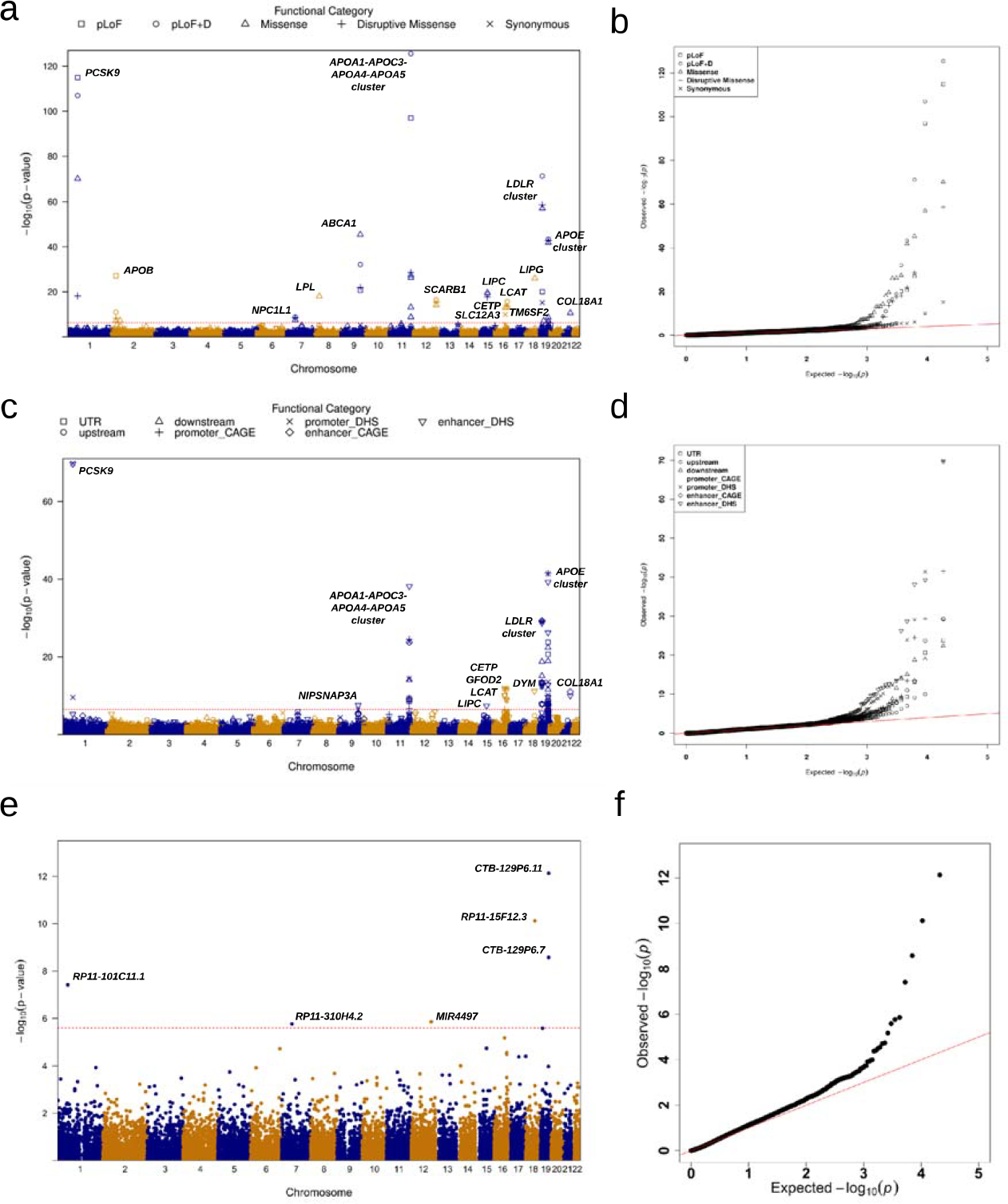
Manhattan plots and Q-Q plots for unconditional gene-centric coding, noncoding and ncRNA analysis of low-density lipoprotein cholesterol (LDL-C), high-density lipoprotein cholesterol (HDL-C) and triglycerids (TG) using TOPMed data (n = 61,838). **a**, Manhattan plots for unconditional gene-centric coding analysis of protein-coding gene. The horizontal line indicates a genome-wide MultiSTAAR-O *P* value threshold of 5.00 × 10^−7^. The significant threshold is defined by multiple comparisons using the Bonferroni correction (0.05/(20,000× 5) =5.00 ×10^−7^). Different symbols represent the MultiSTAAR-O *P* value of the protein-coding gene using different functional categories (putative loss-of-function, putative loss-of-function and disruptive missense, missense, disruptive missense, synonymous). **b**, Quantile-quantile plots for unconditional gene-centric coding analysis of protein-coding gene. Different symbols represent the MultiSTAAR-O *P*-value of the gene using different functional categories. **c**, Manhattan plots for unconditional gene-centric noncoding analysis of protein-coding gene. The horizontal line indicates a genome-wide MultiSTAAR-O *P* value threshold of 3.57 × 10^−7^. The significant threshold is defined by multiple comparisons using the Bonferroni correction (0.05/(20,000× 7) =3.57 ×10^−7^). Different symbols represent the MultiSTAAR-O *P* value of the protein-coding gene using different functional categories (upstream, downstream, UTR, promoter_CAGE, promoter_DHS, enhancer_CAGE, enhancer_DHS). Promoter_CAGE and promoter_DHS are the promoters with overlap of Cap Analysis of Gene Expression (CAGE) sites and DNase hypersensitivity (DHS) sites for a given gene, respectively. Enhancer_CAGE and enhancer_DHS are the enhancers in GeneHancer predicted regions with the overlap of CAGE sites and (DHS) sites for a given gene, respectively. **d**, Quantile-quantile plots for unconditional gene-centric noncoding analysis of protein-coding gene. Different symbols represent the MultiSTAAR-O *P*-value of the gene using different functional categories. **e**, Manhattan plots for unconditional gene-centric noncoding analysis of ncRNA gene. The horizontal line indicates a genome-wide MultiSTAAR-O *P* value threshold of 2.50 ×10-^6^. The significant threshold is defined by multiple comparisons using the Bonferroni correction (0.05/(20,000 =2.50 ×10^−^6). **f**, Quantile-quantile plots for unconditional gene-centric noncoding analysis of ncRNA gene. In panels, **a, c** and **e**, the chromosome number are indicated by the colors of dots. In all panels, MultiSTAAR-O is a two-sided test.

**Extended Data Fig. 2.**
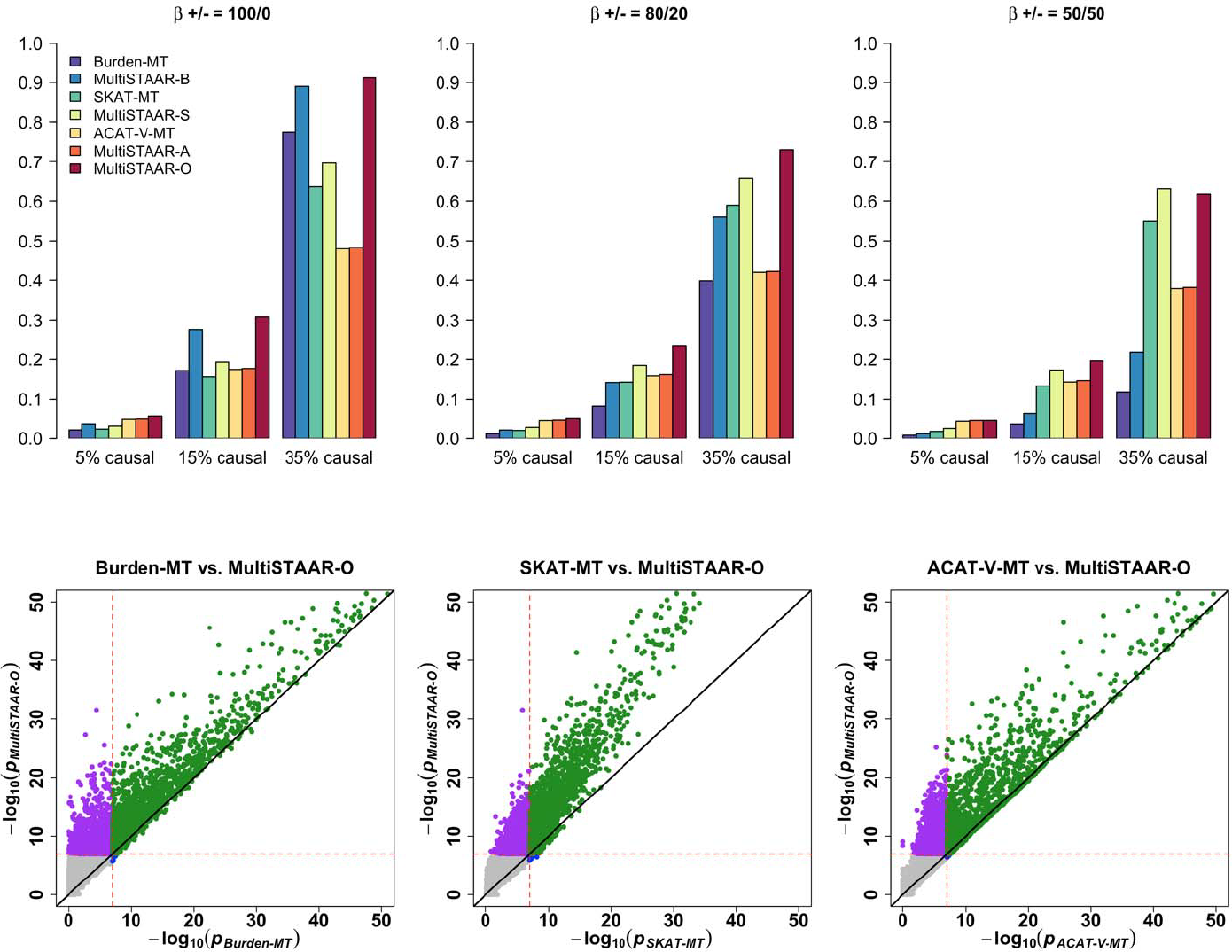
Power comparisons of Burden-MT, SKAT-MT, ACAT-V-MT (MT is short for Multi-Trait) and MultiSTAAR methods when variants in the signal region are associated with one phenotype. Multi-trait Burden, SKAT and ACAT-V tests implemented in MultiSTAAR are denoted by Burden-MT, SKAT-MT and ACAT-V-MT. MultiSTAAR methods incorporating ten functional annotations are denoted by MultiSTAAR-B, MultiSTAAR-S, MultiSTAAR-A and MultiSTAAR-O. In each simulation replicate, a 5-kb region was randomly selected as the signal region. Within each signal region, variants were randomly generated to be causal based on the multivariate logistic model and on average there were 5%, 15% or 35% causal variants in the signal region. The effect sizes of causal variants were *β*_*j*_ = *c*_0_ | log_10_ *MAF*_*j*_|, where *c*_0_ was set to be 0.13. The barplot of power in the top panel consider settings in which the effect sizes for the causal variants are 100% positive (0% negative), 80% positive (20% negative), and 50% positive (50% negative). The scatterplot of *P* values in the bottom panel compare MultiSTAAR-O to Burden-MT, SKAT-MT and ACAT-V-MT when 15% of variants in the signal region are causal variants with all positive effect sizes. Power was estimated as the proportion of the *P* values less than *α* = 10^−7^based on 10^−4^ replicates. Burden-MT, SKAT-MT, ACAT-V-MT, MultiSTAAR-B, MultiSTAAR-S, MultiSTAAR-A and MultiSTAAR-O are two-sided tests. Total sample size considered was 10,000.

**Extended Data Fig. 3.**
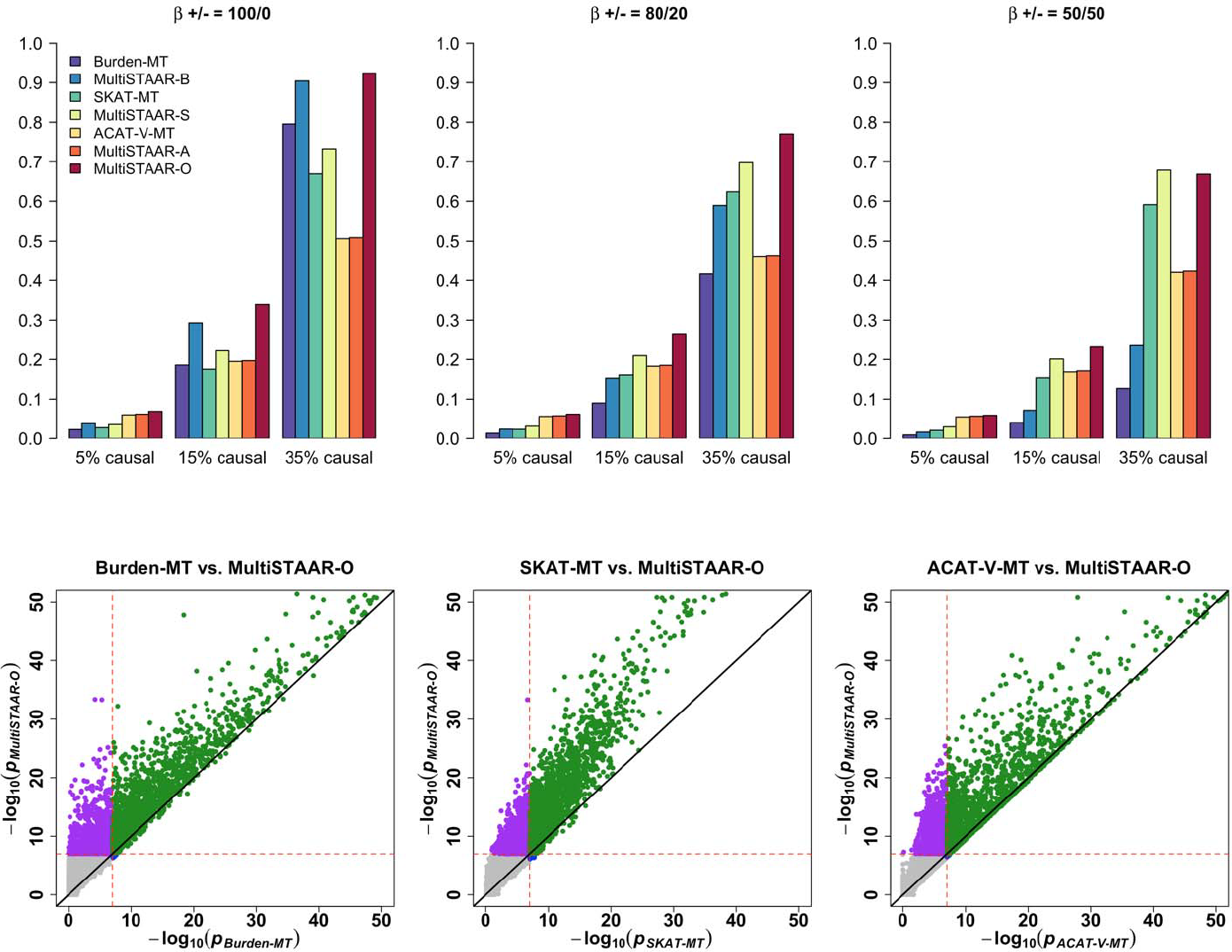
Power comparisons of Burden-MT, SKAT-MT, ACAT-V-MT (MT is short for Multi-Trait) and MultiSTAAR methods when variants in the signal region are associated with two positively correlated phenotypes. In each simulation replicate, a 5-kb region was randomly selected as the signal region. Within each signal region, variants were randomly generated to be causal based on the multivariate logistic model and on average there were 5%, 15% or 35% causal variants in the signal region. The effect sizes of causal variants were *β*_*j*_ = *c*_0_ | log_10_ *MAF*_*j*_|, where *c*0 was set to be 0.1. The barplot of power in the top panel consider settings in which the effect sizes for the causal variants are 100% positive (0% negative), 80% positive (20% negative), and 50% positive (50% negative). The scatterplot of *P* values in the bottom panel compare MultiSTAAR-O to Burden-MT, SKAT-MT and ACAT-V-MT when 15% of variants in the signal region are causal variants with all positive effect sizes. Power was estimated as the proportion of the *P* values less than *α*=10^−7^ based on 10^−4^ replicates. Burden-MT, SKAT-MT, ACAT-V-MT, MultiSTAAR-B, MultiSTAAR-S, MultiSTAAR-A and MultiSTAAR-O are two-sided tests. Total sample size considered was 10,000.

**Extended Data Fig. 4.**
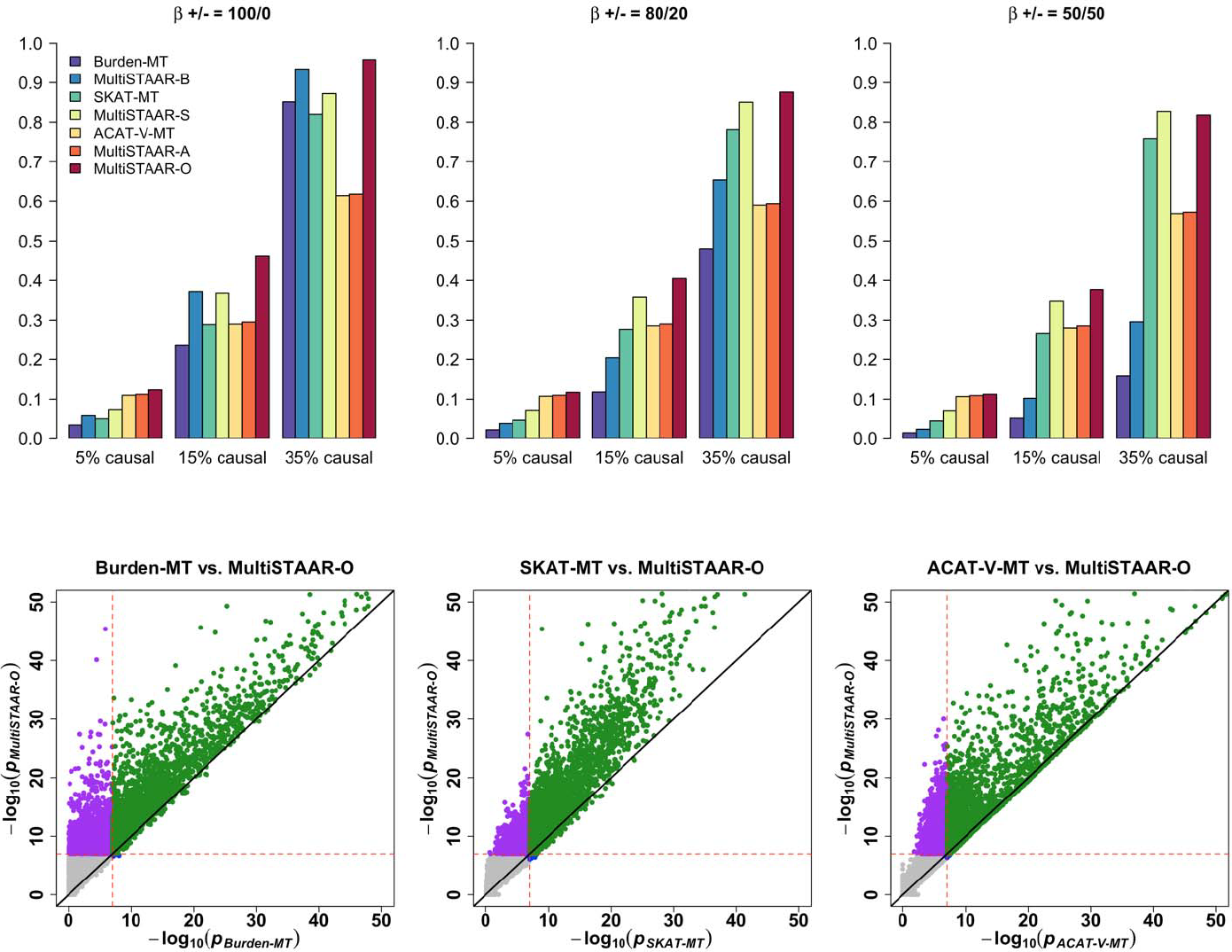
Power comparisons of Burden-MT, SKAT-MT, ACAT-V-MT (MT is short for Multi-Trait) and MultiSTAAR methods when variants in the signal region are associated with two negatively correlated phenotypes. In each simulation replicate, a 5-kb region was randomly selected as the signal region. Within each signal region, variants were randomly generated to be causal based on the multivariate logistic model and on average there were 5%, 15% or 35% causal variants in the signal region. The effect sizes of causal variants were *β*_*j*_ = *c*_0_ | log_10_ *MAF*_*j*_|, where *c*0was set to be 0.1. The barplot of power in the top panel consider settings in which the effect sizes for the causal variants are 100% positive (0% negative), 80% positive (20% negative), and 50% positive (50% negative). The scatterplot of *P* values in the bottom panel compare MultiSTAAR-O to Burden-MT, SKAT-MT and ACAT-V-MT when 15% of variants in the signal region are causal variants with all positive effect sizes. Power was estimated as the proportion of the *P* values less than ^10−7^ based on 10^−4^ replicates. Burden-MT, SKAT-MT, ACAT-V-MT, MultiSTAAR-B, MultiSTAAR-S, MultiSTAAR-A and MultiSTAAR-O are two-sided tests. Total sample size considered was 10,000.

**Extended Data Fig. 5.**
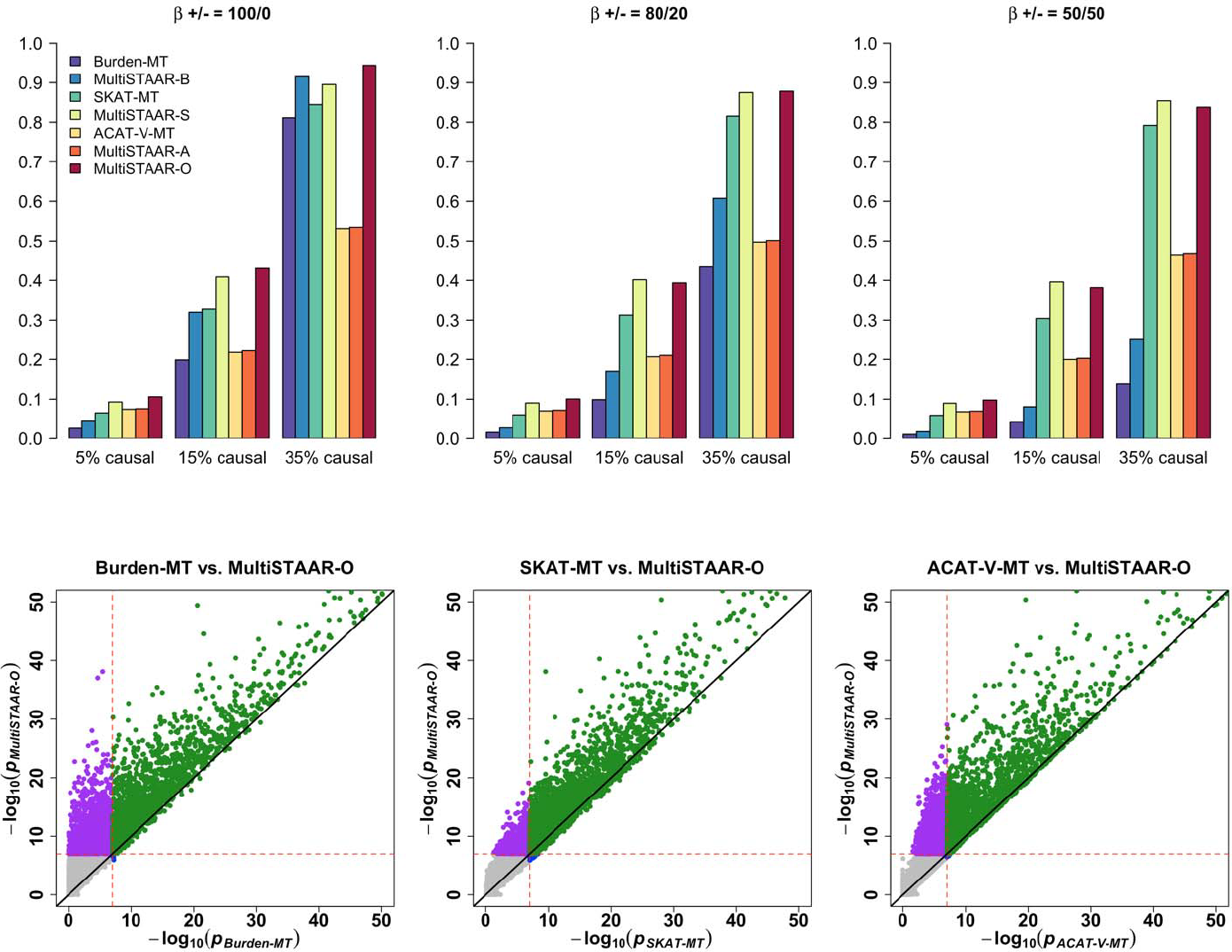
Power comparisons of Burden-MT, SKAT-MT, ACAT-V-MT (MT is short for Multi-Trait) and MultiSTAAR methods when variants in the signal region are associated with three phenotypes. In each simulation replicate, a 5-kb region was randomly selected as the signal region. Within each signal region, variants were randomly generated to be causal based on the multivariate logistic model and on average there were 5%, 15% or 35% causal variants in the signal region. The effect sizes of causal variants were *β*_*j*_ = *c*_0_ | log_10_ *MAF*_*j*_|, where *c*_0_ was set to be 0.07. The barplot of power in the top panel consider settings in which the effect sizes for the causal variants are 100% positive (0% negative), 80% positive (20% negative), and 50% positive (50% negative). The scatterplot of *P* values in the bottom panel compare MultiSTAAR-O to Burden-MT, SKAT-MT and ACAT-V-MT when 15% of variants in the signal region are causal variants with all positive effect sizes. Power was estimated as the proportion of the *P* values less than 10^−7^ based on 10^−4^ replicates. Burden-MT,SKAT-MT, ACAT-V-MT, MultiSTAAR-B, MultiSTAAR-S, MultiSTAAR-A and MultiSTAAR-O are two-sided tests. Total sample size considered was 10,000.

## Methods

### Ethics statement

This study relied on analyses of genetic data from TOPMed cohorts. The study has been approved by the TOPMed Publications Committee, TOPMed Lipids Working Group and all the participating cohorts, including Old Order Amish (phs000956.v1.p1), Atherosclerosis Risk in Communities Study (phs001211), Mt Sinai BioMe Biobank (phs001644), Coronary Artery Risk Development in Young Adults (phs001612), Cleveland Family Study (phs000954), Cardiovascular Health Study (phs001368), Diabetes Heart Study (phs001412), Framingham Heart Study (phs000974), Genetic Study of Atherosclerosis Risk (phs001218), Genetic Epidemiology Network of Arteriopathy (phs001345), Genetic Epidemiology Network of Salt Sensitivity (phs001217), Genetics of Lipid Lowering Drugs and Diet Network (phs001359), Hispanic Community Health Study - Study of Latinos (phs001395), Hypertension Genetic Epidemiology Network and Genetic Epidemiology Network of Arteriopathy (phs001293), Jackson Heart Study (phs000964), Multi-Ethnic Study of Atherosclerosis (phs001416), San Antonio Family Heart Study (phs001215), Genome-wide Association Study of Adiposity in Samoans (phs000972), Taiwan Study of Hypertension using Rare Variants (phs001387), and Women’s Health Initiative (phs001237), where the accession numbers are provided in parenthesis. The use of human genetics data from TOPMed cohorts was approved by the Harvard T.H. Chan School of Public Health IRB (IRB13-0353).

### Notation and model

Suppose there are *n* subjects with a total of *M* variants sequenced across the whole genome. For the *i*-th subject, let ***Y***_*i*_ = (*Y*_*i*1_, *Y*_*i*2_,… *Y*_*iK*_)^*T*^ denote a vector of *K* quantitative phenotypes; ***X***_*i*_ = (*X*_*i*1_,*X*_*i*2_,…*X*_*iq*_)^*T*^ denotes *q* covariates, such as age, gender and ancestral principal components; ***G***_*i*_ = (*G*_*i*1_,*G*_*i*2_,…*G*_*ip*_)^*T*^ denotes the genotype matrix of the *p* genetic variants in a variant set. Since these *K* phenotypes may be defined on different measurement scales, we assume that each phenotype has been rescaled to have zero mean and unit variance.

When the data consist of unrelated samples, we consider the following Multivariate Linear Model (MLM)

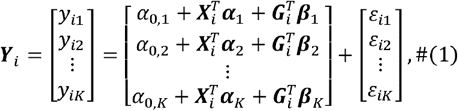

where *α*_*0k*_, is an intercept, ***α*** _*k*_ = (*α* _1,*k*_, *α* _2,*k*_,… *α* _*q,k*_)^*T*^ and ***β*** _*k*_ = (*β* _1,*k*_, *β*_2,*k*_,… *β*_*p,k*_)^*T*^ are column vectors of regression coefficients for covariates ***X****i* and genotype ***G****i* in phenotype *k*, respectively. The error terms ***ε***_*i*_= (*ε*_*i*1,_ *ε*_*i*2,_ *ε*_*iK*_)^*T*^ are independent and identically distributed and follow a multivariate normal distribution with mean a vector of zeros and variance-covariance matrix **Σ***K*x*K*, assumed identical for all subjects. For all *n* subjects, using matrix notation we can write model (1) as

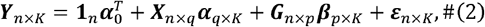

Where **1**_*n*_ is a column vector of 1’s with length *n*, ***α*** _0_ = (*α* _*0*,1_, *α* _*0*,2_,… *α* _*0,K*_)^*T*^ is a column vector of regression intercepts, the *k*-th columns of *α* _*q*x*K*_ and *β* _*p*x*K*_ are *α*_*k*_ and *α*_*k*_, respectively, and ***ε***_*n*_x_*K*_=(*ε*_1,_*ε*_2,…,_*ε*_*n*_)^*T*^ ∼ MatrixNormal_*n,k*_ (**0**_*n*×*k*,_ **I**_*n*×*n*,_ **Σ** _*K*_x_*K*_ **follows** a matrix normal distribution. We calculate the scaled residual for each subject on each phenotype, defined as 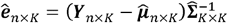, where 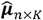 (a matrix of fitted values) and 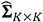 are estimated under the null MLM 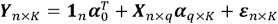, where no variant has any effect on any outcome.

When the data consist of related samples, we consider the following Multivariate Linear Mixed Model (MLMM)^19,47,48^

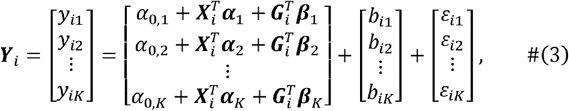

where the random effects *b*_*ik*_ account for relatedness and remaining population structure unaccounted by ancestral PCs^20^. We assume that ***b***_*n*_x_*K*_=(*b*_*ik*_)_*n*×*k*_ ∼ MatrixNormal_*n,k*_ (**0**_*n*×*k*,_ **Φ**_*n*×*n*,_ **Θ**_*K*x*K*_) with a variance component matrix **Θ**_*K*x*K*_ and a sparse genetic relatedness matrix **Φ** _*n*×*n*_^21,22^. For all *n* subjects, using matrix notation we can rewrite equation (3) as

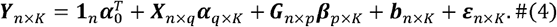

We calculate the scaled residual for each subject on each phenotype, defined as

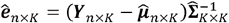, where 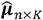 and 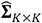 are estimated under the null MLMM 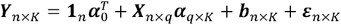 Under both MLM and MLMM, our goal is to test for an association between a set of *p* genetic variants and *K* quantitative phenotypes, adjusting for covariates and relatedness. This corresponds to testing H_0_: β_*1*_ =β_*1*_=…β_*k*_*=0*.

### Multi-trait rare variant association tests using MultiSTAAR

Single-trait score-based aggregation methods^5-9^ can be extended to allow for jointly testing the association between rare variants in a variant set and multiple quantitative phenotypes. For a given variant set, let 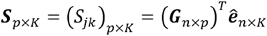 denote the matrix of score statistics where *S*_*jk*_ is the score statistic for the *j*-th variant on the *k*-th phenotype. For multi-trait burden test using MultiSTAAR (Burden-MT), we consider test statistic

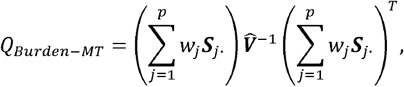

where *w*_*j*_ is the weight defined as a function of the MAF for the *j* -th variant^4,18^,***S****j*· =(*S*_*j*1,_ *S*_*j*2_,…, *S*_*j*K_) is the *j*-th row of ***S*** and 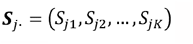 is the *j*-th row of ***S*** and 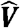 is the estimated variance-covariance matrix of 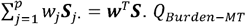 asymptotically follows a standard chi-square distribution with *K* degrees of freedom under the null hypothesis, and its *P* value can be obtained analytically while accounting for LD between variants and correlation between phenotypes.

For multi-trait SKAT using MultiSTAAR (SKAT-MT), we consider the statistic

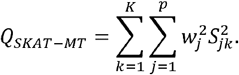

***Q***_*SKAT-MT*_ asymptotically follows a mixture of chi-square distributions under the null hypothesis, and its *P* value can be obtained analytically while accounting for LD between variants and correlation between phenotypes^14,15^.

For multi-trait ACAT-V using MultiSTAAR (ACAT-V-MT), we propose test statistic

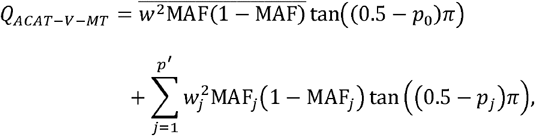

where *p*^′^ is the number of variants with a minor allele count (MAC) greater than 10 and *p*_*j*_ is the multi-trait association *P* value of individual variant *j* for those variants with a MAC>10, whose test statistic is given by the *K* degrees of freedom multivariate score test

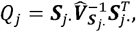

where 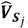 is the estimated variance-covariance matrix of ***S***_***j***·_ ;*p*_0_ is the multi-trait burden test *P* value of extremely rare variants with an MAC ≤10 as described above and 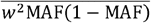 is the average of the weights 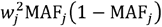 among the extremely rare variants with an MAC≤10.*Q*_*ACAT*-*V*-*MT*_ is approximated well by a scaled Cauchy distribution under the null hypothesis, and its *P* value can be obtained analytically while accounting for LD between variants and correlation between phenotypes^9,49^. Note that when *K*=1, the multi-trait burden, SKAT, and ACAT-V tests reduce to the original single-trait burden, SKAT and ACAT-V tests.

Suppose we have a collection of *L* annotations, let *A*_*jl*_ denote the *l*-th annotation for the *j*th variant in the variant set. We define the functionally-informed multi-trait burden, SKAT and ACAT-V test statistics weighted by the *l*-th annotation as follows

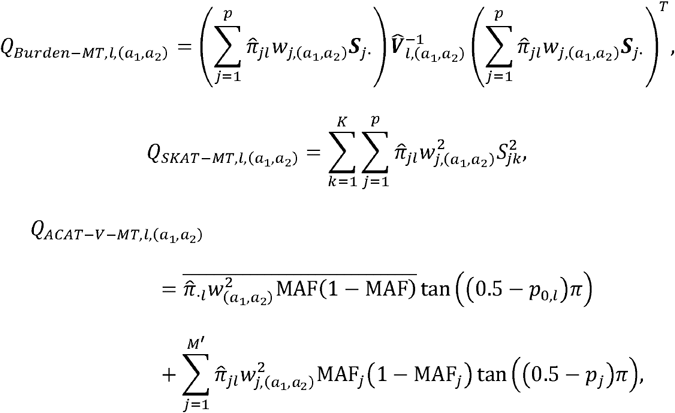

where 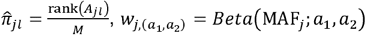 with 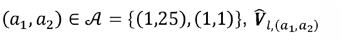 is the estimated variance-covariance matrix of 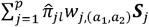, and 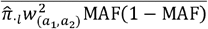 is the average of the weights 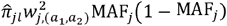 among the extremely rare variants with MAC≤10. Finally, we define the omnibus MultiSTAAR-O test statistic as

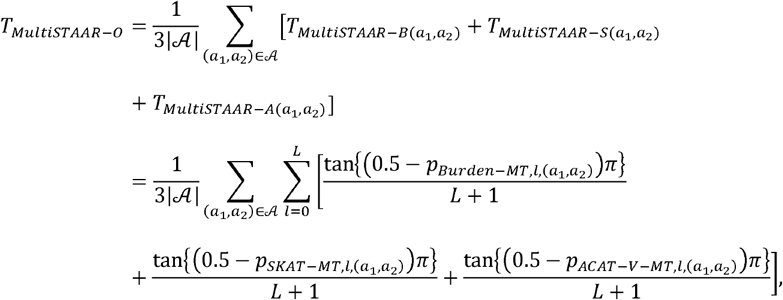

and the *P* value of *T*_*MultiSTAAR-0*_ can be calculated by

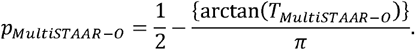

## Data simulation

### Type I error rate simulations

We performed simulation studies to evaluate how accurately MultiSTAAR controls the type I error rate. We generated three quantitative traits from a multivariate linear model, conditional on two covariates

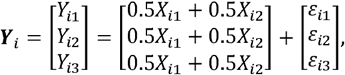

where *X*_*i*1_ ∼ *N* (*0,1), X*_*i*2_ ∼ Bernoulli(0.5) and

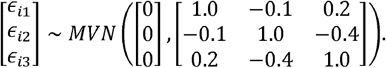

The correlation matrix of error terms *ε*_*i*_= (*ε*_*i*1,_ *ε*_*i*2,_ *ε*_*i*3_)^*T*^ was chosen to mimic the correlations between three lipid traits LDL-C, HDL-C and TG, estimated from the TOPMed data^26^. We considered a sample size of 10,000 and generated genotypes by simulating 20,000 sequences for 100 different regions each spanning 1 Mb. The data generation used the calibration coalescent model (COSI)^29^ with parameters set to mimic the LD structure of African Americans. In each simulation replicate, 10 annotations were generated as *A*_1_,…, *A*_10_ all independently and identically distributed as *N*(0,1)for each variant, and we randomly selected 5-kb regions from these 1-Mb regions for type I error rate simulations. We applied MultiSTAAR-B, MultiSTAAR-S, MultiSTAAR-A and MultiSTAAR-O by incorporating MAFs and the 10 annotations together with Burden-MT, SKAT-MT and ACAT-V-MT tests. We repeated the procedure with 10^8^ replicates to examine the type I error rate at levels *β=*10^−4^, 10^−5^ .and 10^−6^.

### Empirical power simulations

Next, we carried out simulation studies under a variety of configurations to assess the the power of MultiSTAAR-O, and how its incorporation of multiple functional annotations affects power compared to the multi-trait burden, SKAT, and ACAT-V tests implemented in MultiSTAAR. In each simulation replicate, we randomly selected 5-kb regions from a 1-Mb region for power evaluations. For each selected 5-kb region, we generated three quantitative traits from a multivariate linear model

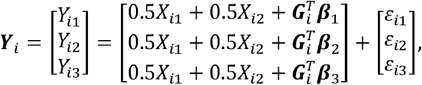

where *X*_*i*1,_ *X*_*i*1_, *ε*_*i*_ were defined as in the type I error rate simulations, ***G***i = (*G*_*i*1_,*G*_*i*2_,…*G*_*ip*_)^*T*^ and ***β =*** *(β*_1,*k*_,*β*_2,*k*,_ *β*_*p,k*_)*>*)^*T*^ were the genotypes and effect sizes of the *p* genetic variants in the signal region.

The genetic effect of variant *j* on phenotype *k* was defined as β _*j,k*_ = *c*_*j*_*d*_*k*_ *γ*_*j*_ to allow for heterogeneous effect sizes among variants and phenotypes. Specifically, we generated the causal variant indicator J according to a logistic model

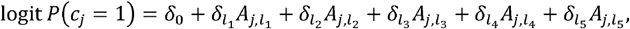

where {*l*_1_,…,*l*_5_} ⊂ {1,…,10} were randomly sampled for each region. For different regions, causality of variants depended on different sets of annotations. We set δ_*l =*_ log(5) for all annotations and varied the proportions of causal variants in the signal region by setting δ_0 *=*_ logit (0.0015), logit (0.0015) and logit (0.18) which corresponds to averaging 5%, 15% and 35% causal variants in the signal region, respectively. We considered four scenarios of phenotypic indicator *d*_*k*_ that reflect different underlying genetic architectures across phenotypes: (*d*_1,_ *d*_2,_ *d*_3_) = (1,0,0), (1,0,1), (1,1,0) and(1,1,1). These correspond to causal variants in the signal region being associated with (1) one phenotype only, (2) two positively correlated phenotypes, (3) two negatively correlated phenotypes and (4) all three phenotypes. We modeled the absolute effect sizes of causal variants using | *γ* _*j*_|= *c*_0_ | log_10_ *MAF*_*j*_|, such that it was a decreasing function of MAF.*c*_0_ was set to be 0.13, 0.1, 0.1 and 0.07, respectively, to ensure a decent power of tests under each scenario. We additionally varied the proportions of causal variant effect size directions (signs of *r*_*j*_) by randomly generating 100%, 80%, and 50% variants on average to have positive effects. We applied MultiSTAAR-B, MultiSTAAR-S, MultiSTAAR-A, and MultiSTAAR-O using MAFs and all 10 annotations together with Burden-MT, SKAT-MT and ACAT-V-MT tests. We repeated the procedure with 10^4^ replicates to examine the power at level *α* = 10^−7^. The sample size was 10,000 across all scenarios.

### Lipid Traits

Conventionally measured plasma lipids, including LDL-C, HDL-C, and triglycerides, were included for analysis. LDL-C was either calculated by the Friedewald equation when triglycerides were <400 mg/dl or directly measured. Given the average effect of statins, when statins were present, LDL-C was adjusted by dividing by 0.7. Triglycerides were natural log transformed for analysis. Phenotypes were harmonized by each cohort and deposited into the dbGaP TOPMed Exchange Area.

### Multi-trait analysis of lipid levels in the TOPMed WGS data

The TOPMed WGS data consist of multi-ethnic related samples^1^. Race/ethnicity was defined using a combination of self-reported race/ethnicity from participant questionnaires and study recruitment information (**Supplementary Note**)^31^. In this study, we applied MultiSTAAR to perform multi-trait rare variant analysis of three quantitative lipid traits (LDL-C, HDL-C and TG) using 20 study cohorts from the TOPMed Freeze 8 WGS data. LDL-C was adjusted for the presence of medications as before^30^. For each study, we first fit a linear regression model adjusting for age, age^2^, sex for each race/ethnicity-specific group. In addition, for Old Order Amish (OOA), we also adjusted for *APOB* p.R3527Q in LDL-C and TC analyses and adjusted for *APOC3* p.R19Ter in TG and HDL-C analyses^30^.

We performed rank-based inverse normal transformation of the residuals of LDL-C, HDL-C and TG within each race/ethnicity-specific group. We then fit a multivariate linear mixed model for the rank normalized residuals, adjusting for 11 ancestral principal components, ethnicity group indicators, and a variance component for empirically derived sparse kinship matrix to account for population structure, relatedness and correlation between phenotypes.

We next applied MultiSTAAR-O to perform multi-trait variant set analyses for rare variants (MAF < 1%) by scanning the genome, including gene-centric analysis of each protein-coding gene using five coding variant functional categories (putative loss-of-function rare variants, missense rare variants, disruptive missense rare variants, putative loss-of-function and disruptive missense rare variants and synonymous rare variants); seven noncoding variant functional categories (promoter rare variants overlaid with CAGE sites, promoter rare variants overlaid with DHS sites, enhancer rare variants overlaid with CAGE sites, enhancer rare variants overlaid with DHS sites, UTR rare variants, upstream region rare variants, downstream region rare variants) and rare variants in ncRNA genes; and genetic region analysis using 2-kb sliding windows across the genome with a 1-kb skip length. The WGS multi-trait rare variant analysis was performed using the R packages MultiSTAAR (version 0.9.7, https://github.com/xihaoli/MultiSTAAR) and STAARpipeline (version 0.9.7, https://github.com/xihaoli/STAARpipeline). The WGS rare variant single-trait analysis of LDL-C, HDL-C and TG was performed using the R package STAARpipeline (version 0.9.7, https://github.com/xihaoli/STAARpipeline). Both multi-trait and single-trait analyses results were summarized and visualized using the R package STAARpipelineSummary (version 0.9.7, https://github.com/xihaoli/STAARpipelineSummary).

### Genome build

All genome coordinates are given in NCBI GRCh38/UCSC hg38.

## Statistics and reproducibility

Sample size was not predetermined. The multi-trait analysis consists of 20 study cohorts of TOPMed Freeze 8 and had 61,838 samples with lipid traits. We did not use any study design that required randomization or blinding.

## Data availability

This paper used the TOPMed Freeze 8 WGS data and lipids phenotype data. Genotype and phenotype data are both available in database of Genotypes and Phenotypes. The TOPMed WGS data were from the following twenty study cohorts (accession numbers provided in parentheses): Old Order Amish (phs000956.v1.p1), Atherosclerosis Risk in Communities Study (phs001211), Mt Sinai BioMe Biobank (phs001644), Coronary Artery Risk Development in Young Adults (phs001612), Cleveland Family Study (phs000954), Cardiovascular Health Study (phs001368), Diabetes Heart Study (phs001412), Framingham Heart Study (phs000974), Genetic Study of Atherosclerosis Risk (phs001218), Genetic Epidemiology Network of Arteriopathy (phs001345), Genetic Epidemiology Network of Salt Sensitivity (phs001217), Genetics of Lipid Lowering Drugs and Diet Network (phs001359), Hispanic Community Health Study - Study of Latinos (phs001395), Hypertension Genetic Epidemiology Network and Genetic Epidemiology Network of Arteriopathy (phs001293), Jackson Heart Study (phs000964), Multi-Ethnic Study of Atherosclerosis (phs001416), San Antonio Family Heart Study (phs001215), Genome-wide Association Study of Adiposity in Samoans (phs000972), Taiwan Study of Hypertension using Rare Variants (phs001387), and Women’s Health

Initiative (phs001237). The sample sizes, ancestry and phenotype summary statistics of these cohorts are given in **Supplementary Table 2**.

The functional annotation data are publicly available and were downloaded from the following links: GRCh38 CADD v1.4 (https://cadd.gs.washington.edu/download); ANNOVAR dbNSFP v3.3a (https://annovar.openbioinformatics.org/en/latest/user-guide/download); LINSIGHT (https://github.com/CshlSiepelLab/LINSIGHT); FATHMM-XF (http://fathmm.biocompute.org.uk/fathmm-xf); FANTOM5 CAGE (https://fantom.gsc.riken.jp/5/data); GeneCards (https://www.genecards.org; v4.7 for hg38); and Umap/Bismap (https://bismap.hoffmanlab.org; ‘before March 2020’ version). In addition, recombination rate and nucleotide diversity were obtained from Gazal et al^50^. The whole-genome individual functional annotation data was assembled from a variety of sources and the computed annotation principal components are available at the Functional Annotation of Variant-Online Resource (FAVOR) site (https://favor.genohub.org)^51^ and the FAVOR database (https://doi.org/10.7910/DVN/1VGTJI)^52^.

## Code availability

MultiSTAAR is implemented as an open source R package available at https://github.com/xihaoli/MultiSTAAR and https://content.sph.harvard.edu/xlin/software.html. Data analysis was performed in R (4.1.0). STAAR v0.9.7 and MultiSTAAR v0.9.7 were used in simulation and real data analysis and implemented as open-source R packages available at https://github.com/xihaoli/STAAR and https://github.com/xihaoli/MultiSTAAR. The assembled functional annotation data were downloaded from FAVOR using Wget (https://www.gnu.org/software/wget/wget.html).

## References

1. Taliun, D. et al. Sequencing of 53,831 diverse genomes from the NHLBI TOPMed Program. Nature 590, 290–299 (2021).

2. The “All of Us” Research Program. New England Journal of Medicine 381, 668–676 (2019).

3. Halldorsson, B.V. et al. The sequences of 150,119 genomes in the UK Biobank. Nature 607, 732–740 (2022).

4. Lee, S., Abecasis, Gonçalo R., Boehnke, M. & Lin, X. Rare-Variant Association Analysis: Study Designs and Statistical Tests. The American Journal of Human Genetics 95, 5–23 (2014).

5. Li, B. & Leal, S.M. Methods for Detecting Associations with Rare Variants for Common Diseases: Application to Analysis of Sequence Data. The American Journal of Human Genetics 83, 311–321 (2008).

6. Madsen, B.E. & Browning, S.R. A Groupwise Association Test for Rare Mutations Using a Weighted Sum Statistic. PLOS Genetics 5, e1000384 (2009).

7. Morris, A.P. & Zeggini, E. An evaluation of statistical approaches to rare variant analysis in genetic association studies. Genetic Epidemiology 34, 188–193 (2010).

8. Wu, Michael C. et al. Rare-Variant Association Testing for Sequencing Data with the Sequence Kernel Association Test. The American Journal of Human Genetics 89, 82–93 (2011).

9. Liu, Y. et al. ACAT: A Fast and Powerful p Value Combination Method for Rare-Variant Analysis in Sequencing Studies. The American Journal of Human Genetics 104, 410–421 (2019).

10. Solovieff, N., Cotsapas, C., Lee, P.H., Purcell, S.M. & Smoller, J.W. Pleiotropy in complex traits: challenges and strategies. Nature Reviews Genetics 14, 483–495 (2013).

11. Sivakumaran, S. et al. Abundant Pleiotropy in Human Complex Diseases and Traits. The American Journal of Human Genetics 89, 607–618 (2011).

12. Abdellaoui, A., Yengo, L., Verweij, K.J.H. & Visscher, P.M. 15 years of GWAS discovery: Realizing the promise. The American Journal of Human Genetics (2023).

13. Watanabe, K. et al. A global overview of pleiotropy and genetic architecture in complex traits. Nature Genetics 51, 1339–1348 (2019).

14. Wu, B. & Pankow, J.S. Sequence Kernel Association Test of Multiple Continuous Phenotypes. Genetic Epidemiology 40, 91–100 (2016).

15. Dutta, D., Scott, L., Boehnke, M. & Lee, S. Multi-SKAT: General framework to test for rare-variant association with multiple phenotypes. Genetic Epidemiology 43, 4–23 (2019).

16. Luo, L. et al. Multi-trait analysis of rare-variant association summary statistics using MTAR. Nature Communications 11, 2850 (2020).

17. Broadaway, K.A. et al. A Statistical Approach for Testing Cross-Phenotype Effects of Rare Variants. The American Journal of Human Genetics 98, 525–540 (2016).

18. Li, X. et al. Dynamic incorporation of multiple in silico functional annotations empowers rare variant association analysis of large whole-genome sequencing studies at scale. Nature Genetics 52, 969–983 (2020).

19. Sammel, M., Lin, X. & Ryan, L. Multivariate linear mixed models for multiple outcomes. Statistics in Medicine 18, 2479–2492 (1999).

20. Conomos, M.P., Miller, M.B. & Thornton, T.A. Robust Inference of Population Structure for Ancestry Prediction and Correction of Stratification in the Presence of Relatedness. Genetic Epidemiology 39, 276–293 (2015).

21. Conomos, Matthew P., Reiner, Alexander P., Weir, Bruce S. & Thornton, Timothy A. Model-free Estimation of Recent Genetic Relatedness. The American Journal of Human Genetics 98, 127–148 (2016).

22. Gogarten, S.M. et al. Genetic association testing using the GENESIS R/Bioconductor package. Bioinformatics 35, 5346–5348 (2019).

23. Lee, P.H. et al. Principles and methods of in-silico prioritization of non-coding regulatory variants. Human Genetics 137, 15–30 (2018).

24. Li, Z. et al. A framework for detecting noncoding rare-variant associations of large-scale whole-genome sequencing studies. Nature Methods 19, 1599–1611 (2022).

25. Morrison, A.C. et al. Practical approaches for whole-genome sequence analysis of heart-and blood-related traits. The American Journal of Human Genetics 100, 205–215 (2017).

26. Selvaraj, M.S. et al. Whole genome sequence analysis of blood lipid levels in >66,000 individuals. Nature Communications 13, 5995 (2022).

27. Liu, Z. & Lin, X. Multiple phenotype association tests using summary statistics in genome-wide association studies. Biometrics 74, 165–175 (2018).

28. Teslovich, T.M. et al. Biological, clinical and population relevance of 95 loci for blood lipids. Nature 466, 707 (2010).

29. Schaffner, S.F. et al. Calibrating a coalescent simulation of human genome sequence variation. Genome research 15, 1576–1583 (2005).

30. Natarajan, P. et al. Deep-coverage whole genome sequences and blood lipids among 16,324 individuals. Nature Communications 9, 3391 (2018).

31. Stilp, A.M. et al. A System for Phenotype Harmonization in the National Heart, Lung, and Blood Institute Trans-Omics for Precision Medicine (TOPMed) Program. American Journal of Epidemiology (2021).

32. Frankish, A. et al. GENCODE reference annotation for the human and mouse genomes. Nucleic acids research 47, D766–D773 (2019).

33. Dong, C. et al. Comparison and integration of deleteriousness prediction methods for nonsynonymous SNVs in whole exome sequencing studies. Human Molecular Genetics 24, 2125–2137 (2014).

34. Li, Z. et al. A framework for detecting noncoding rare variant associations of large-scale whole-genome sequencing studies. bioRxiv, 2021.11.05.467531 (2021).

35. Kircher, M. et al. A general framework for estimating the relative pathogenicity of human genetic variants. Nature Genetics 46, 310–315 (2014).

36. Huang, Y.-F., Gulko, B. & Siepel, A. Fast, scalable prediction of deleterious noncoding variants from functional and population genomic data. Nature Genetics 49, 618–624 (2017).

37. Rogers, M.F. et al. FATHMM-XF: accurate prediction of pathogenic point mutations via extended features. Bioinformatics 34, 511–513 (2017).

38. Buniello, A. et al. The NHGRI-EBI GWAS Catalog of published genome-wide association studies, targeted arrays and summary statistics 2019. Nucleic Acids Research 47, D1005–D1012 (2019).

39. Klarin, D. et al. Genetics of blood lipids among ∼300,000 multi-ethnic participants of the Million Veteran Program. Nature Genetics 50, 1514–1523 (2018).

40. Forrest, A.R. et al. A promoter-level mammalian expression atlas. Nature 507, 462 (2014).

41. Abascal, F. et al. Expanded encyclopaedias of DNA elements in the human and mouse genomes. Nature 583, 699–710 (2020).

42. Andersson, R. et al. An atlas of active enhancers across human cell types and tissues. Nature 507, 455–461 (2014).

43. Fishilevich, S. et al. GeneHancer: genome-wide integration of enhancers and target genes in GeneCards. Database 2017(2017).

44. Li, Z. et al. Dynamic Scan Procedure for Detecting Rare-Variant Association Regions in Whole-Genome Sequencing Studies. The American Journal of Human Genetics 104, 802–814 (2019).

45. McCaw, Z.R., Gao, J., Lin, X. & Gronsbell, J. Leveraging a machine learning derived surrogate phenotype to improve power for genome-wide association studies of partially missing phenotypes in population biobanks. bioRxiv, 2022.12.12.520180 (2022).

46. Li, X. et al. Powerful, scalable and resource-efficient meta-analysis of rare variant associations in large whole genome sequencing studies. Nature Genetics 55, 154–164 (2023).

47. Chen, H. et al. Control for Population Structure and Relatedness for Binary Traits in Genetic Association Studies via Logistic Mixed Models. The American Journal of Human Genetics 98, 653–666 (2016).

48. Chen, H. et al. Efficient Variant Set Mixed Model Association Tests for Continuous and Binary Traits in Large-Scale Whole-Genome Sequencing Studies. The American Journal of Human Genetics 104, 260–274 (2019).

49. Liu, Y. & Xie, J. Cauchy combination test: a powerful test with analytic p-value calculation under arbitrary dependency structures. Journal of the American Statistical Association 115, 393–402 (2020).

## References

50. Gazal, S. et al. Linkage disequilibrium–dependent architecture of human complex traits shows action of negative selection. Nature Genetics 49, 1421–1427 (2017).

51. Zhou, H. et al. FAVOR: functional annotation of variants online resource and annotator for variation across the human genome. Nucleic Acids Research 51, D1300–D1311 (2023).

52. Zhou, H., Arapoglou, T., Li, X., Li, Z. & Lin, X. FAVOR Essential Database. V1 Edition (Harvard Dataverse, 2022).

