## Supplementary Figures for "A statistical framework for powerful multi-trait rare variant analysis in large-scale whole-genome sequencing studies"

**Supplementary Figure 1.** **Scatterplot of *P* values comparing MultiSTAAR-O to Burden-MT, SKAT-MT and ACAT-V-MT (MT is short for Multi-Trait) when variants in the signal region are associated with one phenotype.** In each simulation replicate, a 5-kb region was randomly selected as the signal region. Within each signal region, variants were randomly generated to be causal based on the multivariate logistic model and on average there were 5% (top) or 35% (bottom) causal variants in the signal region. The effect sizes of causal variants were $\beta_{j}=c_{0}|\log_{10} MAF_{j}|$, where $c_{0}$ was set to be 0.13. All causal variants had positive effect sizes. Power was estimated as the proportion of the *P* values less than $\alpha={10}^{-7}$ based on ${10}^{4}$ replicates. Burden-MT, SKAT-MT, ACAT-V-MT and MultiSTAAR-O are two-sided tests. Total sample size considered was 10,000.

**5% Causal Variants**

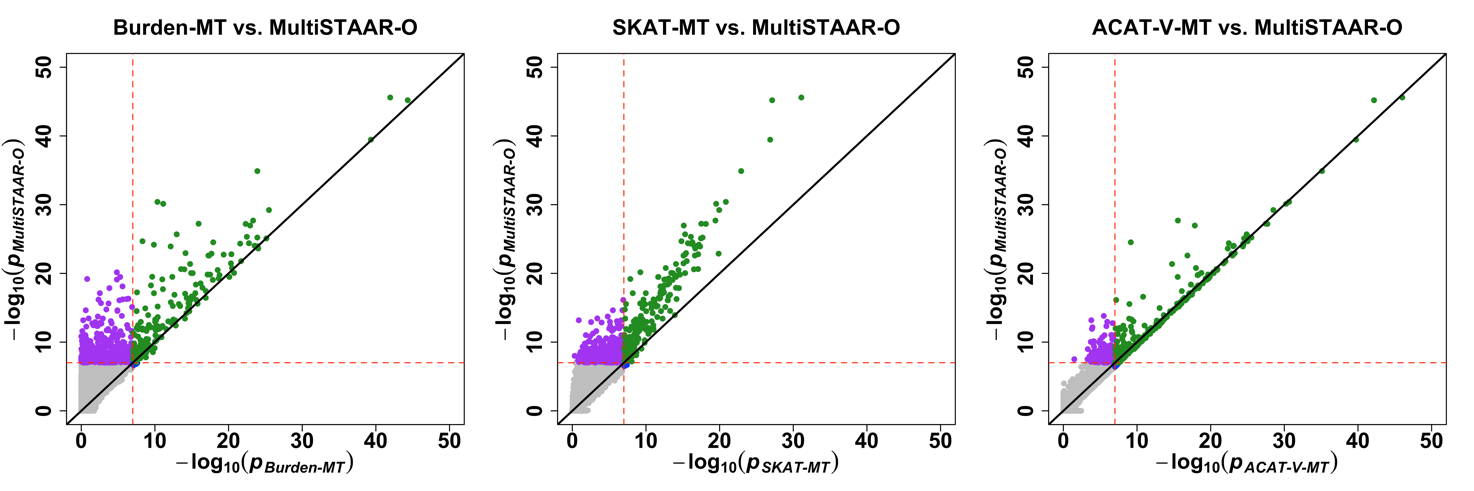

**35% Causal Variants**

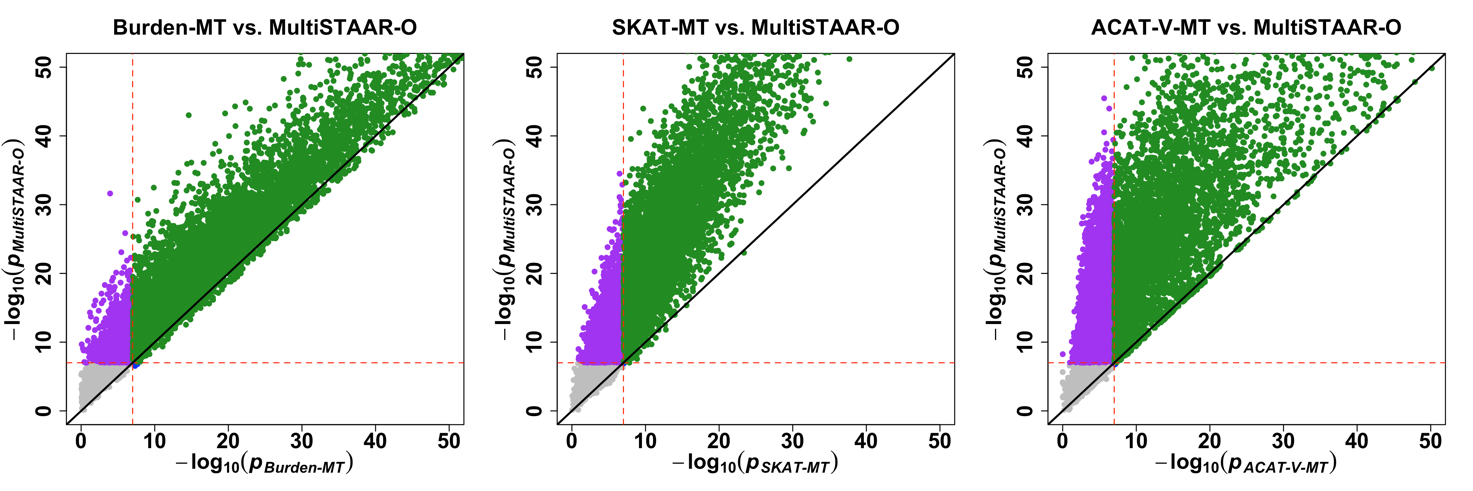

**Supplementary Figure 2.** **Scatterplot of *P* values comparing MultiSTAAR-O to Burden-MT, SKAT-MT and ACAT-V-MT (MT is short for Multi-Trait) when variants in the signal region are associated with two positively correlated phenotypes.** In each simulation replicate, a 5-kb region was randomly selected as the signal region. Within each signal region, variants were randomly generated to be causal based on the multivariate logistic model and on average there were 5% (top) or 35% (bottom) causal variants in the signal region. The effect sizes of causal variants were $\beta_{j}=c_{0}|\log_{10} MAF_{j}|$, where $c_{0}$ was set to be 0.1. All causal variants had positive effect sizes. Power was estimated as the proportion of the *P* values less than $\alpha={10}^{-7}$ based on ${10}^{4}$ replicates. Burden-MT, SKAT-MT, ACAT-V-MT and MultiSTAAR-O are two-sided tests. Total sample size considered was 10,000.

**5% Causal Variants**

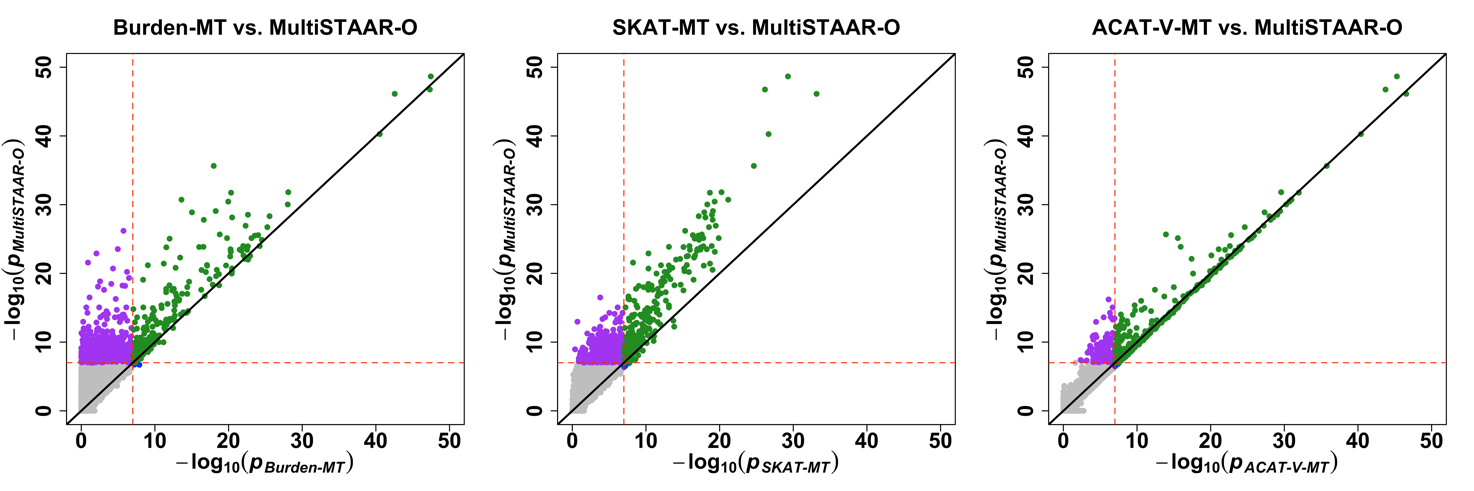

**35% Causal Variants**

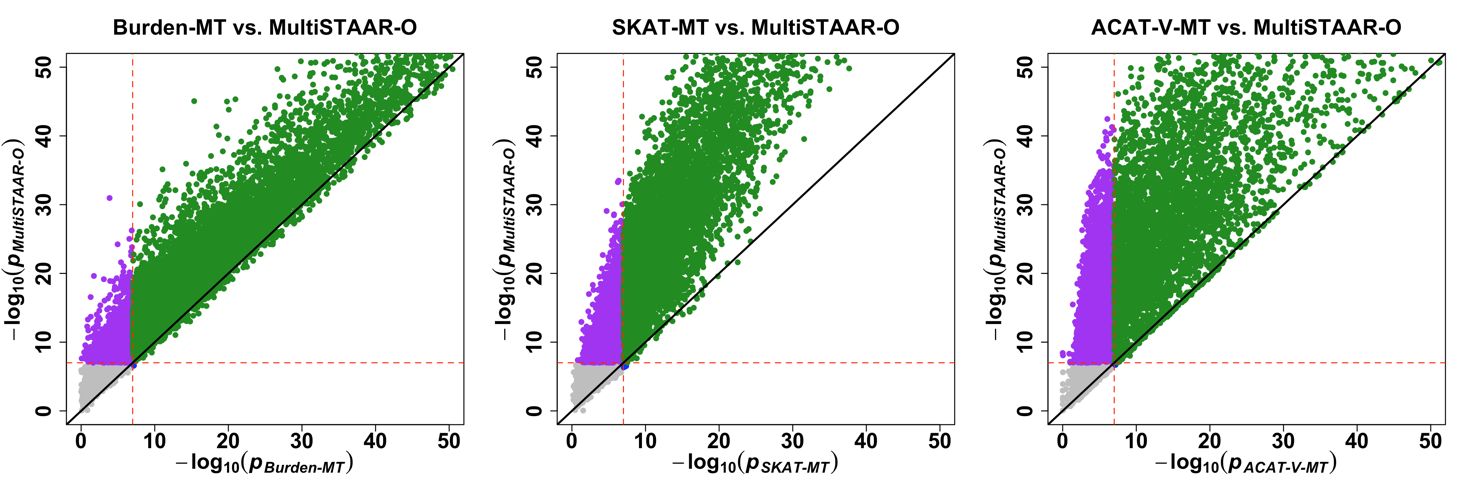

**Supplementary Figure 3.** **Scatterplot of *P* values comparing MultiSTAAR-O to Burden-MT, SKAT-MT and ACAT-V-MT (MT is short for Multi-Trait) when variants in the signal region are associated with two negatively correlated phenotypes.** In each simulation replicate, a 5-kb region was randomly selected as the signal region. Within each signal region, variants were randomly generated to be causal based on the multivariate logistic model and on average there were 5% (top) or 35% (bottom) causal variants in the signal region. The effect sizes of causal variants were $\beta_{j}=c_{0}|\log_{10} MAF_{j}|$, where $c_{0}$ was set to be 0.1. All causal variants had positive effect sizes. Power was estimated as the proportion of the *P* values less than $\alpha={10}^{-7}$ based on ${10}^{4}$ replicates. Burden-MT, SKAT-MT, ACAT-V-MT and MultiSTAAR-O are two-sided tests. Total sample size considered was 10,000.

**5% Causal Variants**

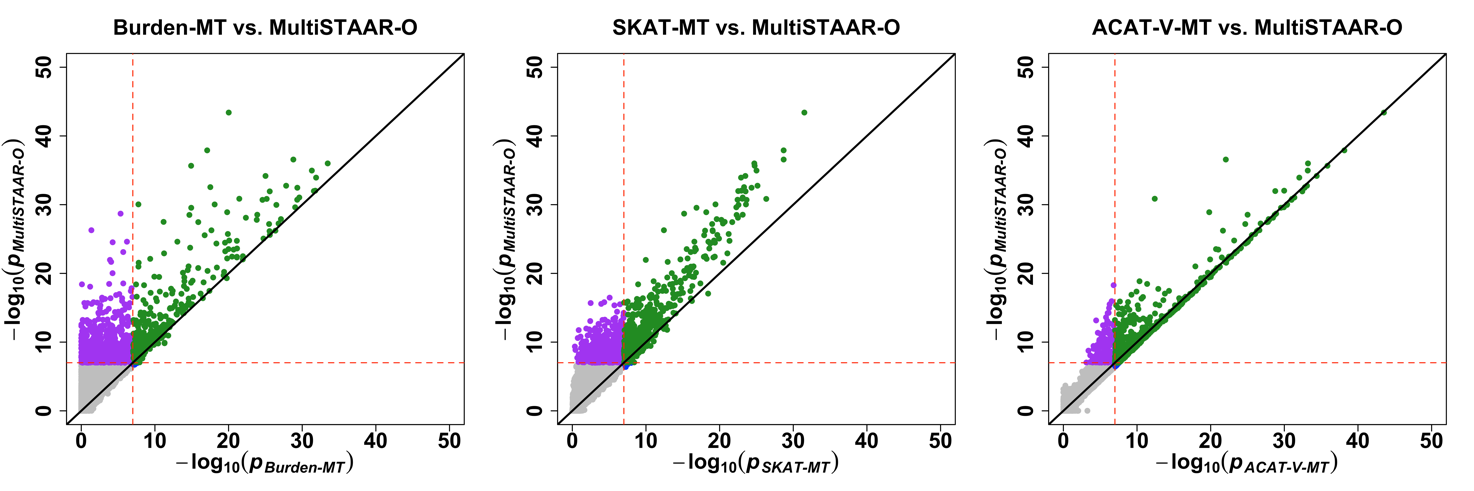

**35% Causal Variants**

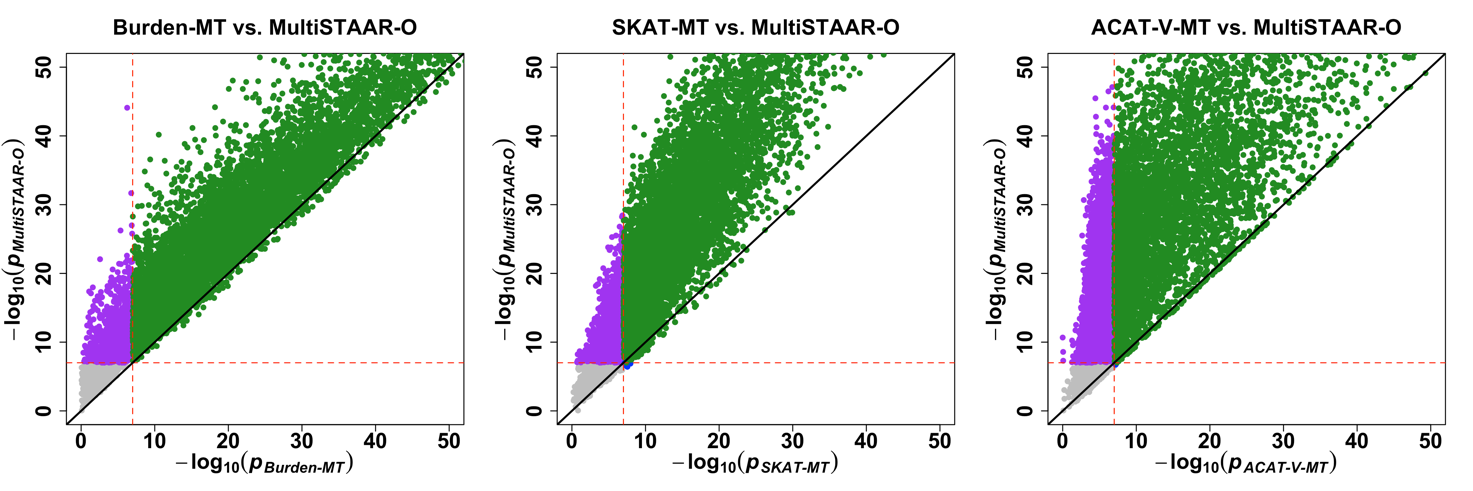

**Supplementary Figure 4.** **Scatterplot of *P* values comparing MultiSTAAR-O to Burden-MT, SKAT-MT and ACAT-V-MT (MT is short for Multi-Trait) when variants in the signal region are associated with three phenotypes.** In each simulation replicate, a 5-kb region was randomly selected as the signal region. Within each signal region, variants were randomly generated to be causal based on the multivariate logistic model and on average there were 5% (top) or 35% (bottom) causal variants in the signal region. The effect sizes of causal variants were $\beta_{j}=c_{0}|\log_{10} MAF_{j}|$, where $c_{0}$ was set to be 0.07. All causal variants had positive effect sizes. Power was estimated as the proportion of the *P* values less than $\alpha={10}^{-7}$ based on ${10}^{4}$ replicates. Burden-MT, SKAT-MT, ACAT-V-MT and MultiSTAAR-O are two-sided tests. Total sample size considered was 10,000.

**5% Causal Variants**

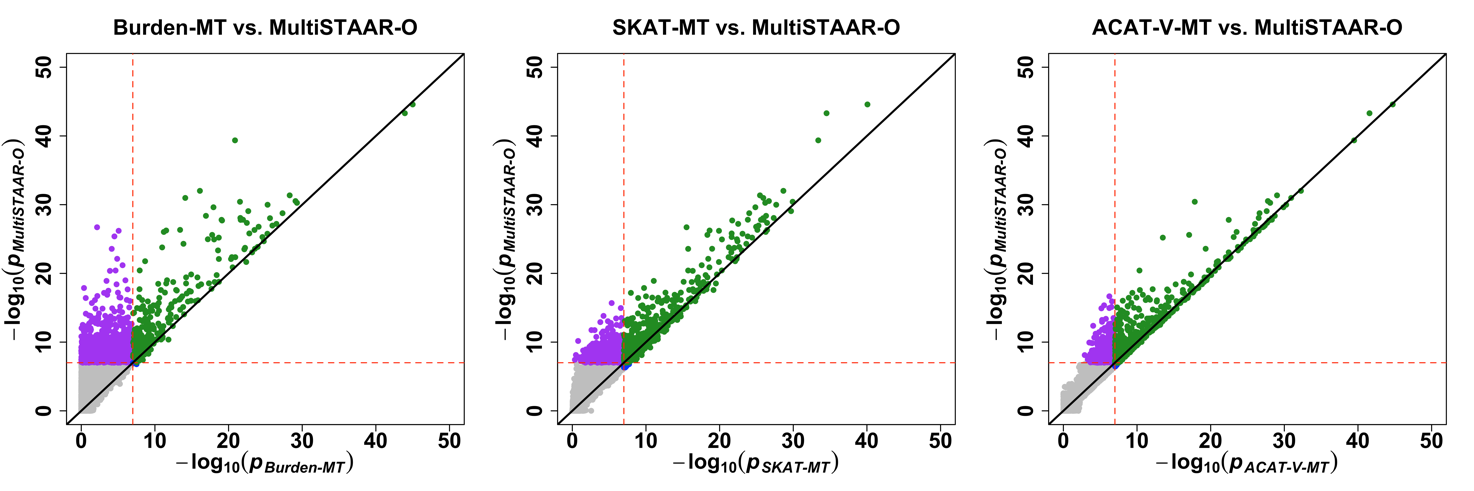

**35% Causal Variants**

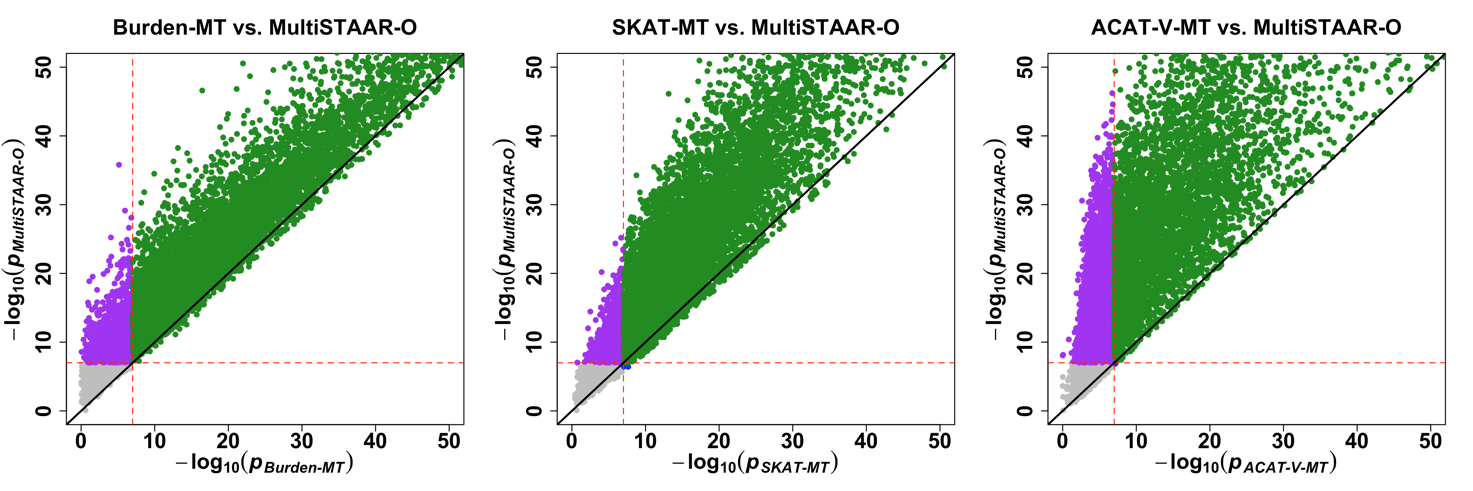
