## Supplementary Note for "A statistical framework for powerful multi-trait rare variant analysis in large-scale whole-genome sequencing studies"

**TOPMed study participants and acknowledgements**

Old Order Amish (OOA)

The Old Order Amish individuals included in this study were participants of several ongoing studies of cardiovascular health carried out at the University of Maryland among relatively healthy volunteers from the Old Order Amish community of Lancaster County, PA and their family members^1,2^.

Whole genome sequencing (WGS) for the Trans-Omics in Precision Medicine (TOPMed) program was supported by the National Heart, Lung and Blood Institute (NHLBI). WGS for “NHLBI TOPMed: Genetics of Cardiometabolic Health in the Amish” (phs000956.v1.p1) was performed at the Broad Institute of MIT and Harvard (3R01HL121007-01S1).

Atherosclerosis Risk in Communities Study (ARIC)

The ARIC study is a population-based prospective cohort study of cardiovascular disease sponsored by the National Heart, Lung, and Blood Institute (NHLBI). ARIC included 15,792 individuals, predominantly European American and African American, aged 45-64 years at baseline (1987-89), chosen by probability sampling from four US communities. Cohort members completed three additional triennial follow-up examinations, a fifth exam in 2011-2013, a sixth exam in 2016-2017, and a seventh exam in 2018-2019. The ARIC study has been described in detail previously^3^.

Whole genome sequencing (WGS) for the Trans-Omics in Precision Medicine (TOPMed) program was supported by the National Heart, Lung and Blood Institute (NHLBI). WGS for “NHLBI TOPMed: Atherosclerosis Risk in Communities (ARIC)” (phs001211) was performed at the Baylor College of Medicine Human Genome Sequencing Center (HHSN268201500015C and 3U54HG003273-12S2) and the Broad Institute of MIT and Harvard (3R01HL092577-06S1). Centralized read mapping and genotype calling, along with variant quality metrics and filtering were provided by the TOPMed Informatics Research Center (3R01HL-117626-02S1; contract HHSN268201800002I). Phenotype harmonization, data management, sample-identity QC, and general study coordination, were provided by the TOPMed Data Coordinating Center (3R01HL-120393-02S1; contract HHSN268201800001I). We gratefully acknowledge the studies and participants who provided biological samples and data for TOPMed.

The Atherosclerosis Risk in Communities study has been funded in whole or in part with Federal funds from the National Heart, Lung, and Blood Institute, National Institutes of Health, Department of Health and Human Services (contract numbers HHSN268201700001I, HHSN268201700002I, HHSN268201700003I, HHSN268201700004I and HHSN268201700005I). The authors thank the staff and participants of the ARIC study for their important contributions.

Mt Sinai BioMe Biobank (BioMe):

The Institute for Personalized Medicine at the Icahn School of Medicine at Mount Sinai

is leading the movement toward diagnosis and classification of disease according to the

patient’s molecular profile. BioMe is the major effort towards this goal, BioMe, an

electronic medical record-linked biobank that enables researchers to conduct genetic,

epidemiologic, molecular, and genomic studies rapidly and efficiently on large

collections of research specimens linked with medical information. BioMed cohort is

composed of White, Black, Hispanic and Asian populations. The Mount Sinai Medical

Center services diverse local communities of upper Manhattan, including Central

Harlem (86% African American), East Harlem (88% Hispanic Latino), and Upper East

Side (88% Caucasian/white) with broad health disparities^4^. Whole genome sequencing

(WGS) for the Trans-Omics in Precision Medicine (TOPMed) program was supported by

the National Heart, Lung and Blood Institute (NHLBI). The Mount Sinai BioMe Biobank

has been supported by The Andrea and Charles Bronfman Philanthropies and in part by

Federal funds from the NHLBI and NHGRI (U01HG00638001; U01HG007417;

X01HL134588).

Coronary Artery Risk Development in Young Adults (CARDIA):

The Coronary Artery Risk Development in Young Adults (CARDIA) Study is a study

examining the development and determinants of clinical and subclinical cardiovascular

disease and their risk factors. It began in 1985-6 with a group of 5115 black and white

men and women aged 18-30 years^5^. The participants were selected so that there would be approximately the same number of people in subgroups of race, gender, education (high school or less and more than high school) and age (18-24 and 25-30) in each of 4 centers: Birmingham, AL; Chicago, IL; Minneapolis, MN; and Oakland, CA. These same participants were asked to participate in follow-up examinations during 1987-1988 (Year 2), 1990-1991 (Year 5), 1992-1993 (Year 7), 1995-1996 (Year 10), 2000-2001 (Year 15), 2005-2006 (Year 20), 2010-2011 (Year 25), and 2015-2016 (Year 30).

Cleveland Family Study (CFS)

The CFS is a family-based longitudinal study that includes participants with laboratory diagnosed sleep apnea, their family members and neighborhood control families followed between 1990 and 2006. Four examinations over 16 years provided measurements of sleep apnea with overnight polysomnography, anthropometry, and other related phenotypes, as detailed previously^6,7^. After an overnight fast, blood was collected which was assayed for lipid levels at the University of Vermont Laboratory for Clinical Biochemistry Research. Lipids (triglycerides, HDL cholesterol) from fasted blood serum were measured by enzymatic methods using Centers for Disease Control and Prevention guidelines^8^.

Whole genome sequencing (WGS) for the Trans-Omics in Precision Medicine (TOPMed) program was supported by the National Heart, Lung and Blood Institute (NHLBI). WGS for “NHLBI TOPMed: Cleveland Family Study” (phs000954) was performed at the University of Washington Northwest Genomics Center (3R01HL098433-05S1).

This research was supported by grants HL 046389; HL113338; 1R35HL135818 **from the National Heart, Lung, and Blood Institute (NHLBI)**.

Cardiovascular Health Study (CHS)

The Cardiovascular Health Study is a prospective population-based cohort study of risk factors for CHD and stroke in adults 65 years and older^9^. The main objective is to identify factors related to the onset and course of heart disease and stroke. The four Field Centers are located in Forsyth County, NC; Sacramento County, CA; Washington County, MD; and Pittsburgh, PA. The original cohort of 5201 elderly were recruited in 1989-1990; and in 1992-1993, 687 additional minority participants were recruited and examined. Each community sample was obtained from random samples of the Medicare eligibility lists of the Health Care Financing Administration (HCFA). Eligible to participate were persons living in the household of each sampled individual who were: 1) 65 yr or older; 2) non-institutionalized; 3) expected to remain in the area for 3 yr; and 4) able to give informed consent. Excluded were those wheelchair-bound, receiving hospice care or cancer treatment. The minority cohort was recruited using similar methods. Participants were eligible whether or not they had clinically apparent cardiovascular disease. Subjects were followed with semi-annual contacts, alternating between telephone calls and surveillance clinic visits.

Whole genome sequencing (WGS) for the Trans-Omics in Precision Medicine (TOPMed) program was supported by the National Heart, Lung and Blood Institute (NHLBI). WGS for “NHLBI TOPMed: Cardiovascular Health Study” (phs001368) was performed at the Baylor College of Medicine Human Genome Sequencing Center (HHSN268201500015C). Analyses were limited to those with available DNA who consented to genetic studies.

**This research was supported by contracts HHSN268201200036C, HHSN268200800007C, HHSN268201800001C, N01HC55222, N01HC85079, N01HC85080, N01HC85081, N01HC85082, N01HC85083, N01HC85086, 75N92021D00006, and grants U01HL080295 and U01HL130114 from the National Heart, Lung, and Blood Institute (NHLBI), with additional contribution from the National Institute of Neurological Disorders and Stroke (NINDS). Additional support was provided by R01AG023629 from the National Institute on Aging (NIA). A full list of principal CHS investigators and institutions can be found at** [CHS-NHLBI.org](https://urldefense.proofpoint.com/v2/url?u=https-3A__chs-2Dnhlbi.org&d=DwMGaQ&c=WO-RGvefibhHBZq3fL85hQ&r=b5Nbqbd-i1zGTvpDeTmxllZPAOZcm40GJ2T0r9cfKmI&m=aO3aCp_qn4wQyYNri083AIpCuFbcV5lFdIZtpOTrToo&s=4kzdKhr8oCU2j4wnSvCQB8emkH-NRBqJVoZPcl-ncf4&e=)**.**

Diabetes Heart Study (DHS)

The Diabetes Heart Study (DHS) began as a family-based study enriched for type 2 diabetes (T2D). The initial cohort included 1443 European American and African American participants from 564 families with multiple cases of type 2 diabetes recruited between 1998 and 2006^10^. As an ancillary study, the African American Diabetes Heart Study (AA-DHS) expanded the total number of African Americans to 691 by recruiting additional unrelated participants with type 2 diabetes from 2007 and 2010^11^. All participants were extensively phenotyped for measures of subclinical CVD and other known CVD risk factors. Primary outcomes were quantified burden of vascular calcified plaque in the coronary artery, carotid artery, and abdominal aorta all determined from non-contrast computed tomography scans. For TOPMed, DHS and AA-DHS African American participants with CAC were selected for WGS, prioritizing the inclusion of families.

Whole genome sequencing (WGS) for the Trans-Omics in Precision Medicine (TOPMed) program was supported by the National Heart, Lung and Blood Institute (NHLBI). WGS for “NHLBI TOPMed: Diabetes Heart Study” (phs001412) was performed at the Broad Institute of MIT and Harvard (HHSN268201500014C).

This work was supported by R01 HL92301, R01 HL67348, R01 NS058700, R01 AR48797, R01 DK071891, R01 AG058921, the General Clinical Research Center of the Wake Forest University School of Medicine (M01 RR07122, F32 HL085989), the American Diabetes Association, and a pilot grant from the Claude Pepper Older Americans Independence Center of Wake Forest University Health Sciences (P60 AG10484).

Framingham Heart Study (FHS)

The FHS is a three generational prospective cohort that has been described in detail previously^12^. Individuals were initially recruited in 1948 in Framingham, USA to evaluate cardiovascular disease risk factors. The second generation cohort (5,124 offspring of the original cohort) was recruited between 1971 and 1975^13,14^. The third generation cohort (4,095 grandchildren of the original cohort) was collected between 2002 and 2005. Fasting lipid levels were measured at exam 1 of the Offspring (1971-1975) and third generation (2002-2005) cohorts, using standard LRC protocols.

Whole genome sequencing (WGS) for the Trans-Omics in Precision Medicine (TOPMed) program was supported by the National Heart, Lung and Blood Institute (NHLBI). WGS for “NHLBI TOPMed: Whole Genome Sequencing and Related Phenotypes in the Framingham Heart Study” (phs000974.v1.p1) was performed at the Broad Institute of MIT and Harvard (HHSN268201500014C).

The Framingham Heart Study (FHS) acknowledges the support of contracts NO1-HC-25195, HHSN268201500001I and 75N92019D00031 from the National Heart, Lung and Blood Institute and grant supplement R01 HL092577-06S1 for this research. We also acknowledge the dedication of the FHS study participants without whom this research would not be possible. Dr. Vasan is supported in part by the Evans Medical Foundation and the Jay and Louis Coffman Endowment from the Department of Medicine, Boston University School of Medicine.

Genetic Study of Atherosclerosis Risk (GeneSTAR)

GeneSTAR is an ongoing family-based prospective study designed to determine environmental, phenotypic, and genetic causes of premature cardiovascular disease. GeneSTAR was originally conducted in healthy adult European- and African-American siblings of probands with documented early onset coronary disease under 60 years of age at the time of hospitalization in any of 10 Baltimore area hospitals from 1982-2006. Participants were screened for traditional coronary disease and stroke risk factors and have been followed regularly to ascertain incident cardiovascular disease^15^. Commencing in 2003, the siblings, their offspring, and the coparent of the offspring who were free of cardiovascular disease participated in a 2 week trial of aspirin 81 mg/day with pre and post ex vivo platelet function assessed using multiple agonists and were screened for traditional coronary disease and stroke risk factors^16^. Of the total 3949 participants, 1786 were selected for TOPMed prioritized on complete platelet function measures and largest family size.

Whole genome sequencing (WGS) for the Trans-Omics in Precision Medicine (TOPMed) program was supported by the National Heart, Lung and Blood Institute (NHLBI). WGS for “NHLBI TOPMed: Genetic Study of Atherosclerosis Risk” (phs001218) was performed at Psomagen (formerly Macrogen; 3R01HL112064-04S1), Illumina (R01HL112064), and the Broad Institute of MIT and Harvard (HHSN268201500014C).

GeneSTAR was supported by grants from the National Institutes of Health/National Heart, Lung, and Blood Institute (U01 HL72518, HL087698, HL49762, HL59684, HL58625, HL071025, HL112064), by a grant from the National Institutes of Health/National Institute of Nursing Research (NR0224103), and by a grant from the National Institutes of Health/National Center for Research Resources (M01-RR000052) to the Johns Hopkins General Clinical Research Center.

Genetic Epidemiology Network of Arteriopathy (GENOA)

The Genetic Epidemiology Network of Arteriopathy (GENOA) is one of four networks in the NHLBI Family-Blood Pressure Program (FBPP)^17^. GENOA’s long-term objective is to elucidate the genetics of target organ complications of hypertension, including both atherosclerotic and arteriolosclerotic complications involving the heart, brain, kidneys, and peripheral arteries^18^. The longitudinal GENOA Study recruited European-American and African-American sibships with at least 2 individuals with clinically diagnosed essential hypertension before age 60 years. All other members of the sibship were invited to participate regardless of their hypertension status. Participants were diagnosed with hypertension if they had either 1) a previous clinical diagnosis of hypertension by a physician with current anti-hypertensive treatment, or 2) an average systolic blood pressure ≥ 140 mm Hg or diastolic blood pressure ≥ 90 mm Hg based on the second and third readings at the time of their clinic visit. Only participants of the African-American Cohort were sequenced through TOPMed.

During the first exam (Phase 1; 1996-2000), 1,583 European-Americans from Rochester, MN and 1,854 African-Americans from Jackson, MS were examined. Between 2000 and 2004 (Phase 2), 1,241 participants of the European-American Cohort and 1,482 participants of the African-American cohort returned for a second examination. The second examination of the European-American cohort included computed tomography scans for coronary artery calcification while the second examination of the African-American cohort included an echocardiogram. Between 2009 and 2011, an examination that included computed tomography scans for coronary artery calcification (CAC Study) was conducted on 752 participants of the African-American Cohort.

Every participant with an echocardiogram was selected for whole genome sequencing (WGS) through TOPMed. We then selected 106 African-American participants who had a computed tomography scan for coronary artery calcification but not an echocardiogram or were a sibling of someone already selected for WGS. Finally, we excluded individuals whom we knew were already being whole genome sequenced through TOPMed or another sequencing effort (GENOA participants who overlap with ARIC or JHS participants).

Support for GENOA was provided by the National Heart, Lung and Blood Institute (HL054457, HL054464, HL054481, HL119443, HL085571, and HL087660) of the National Institutes of Health. DNA extraction for “NHLBI TOPMed: Genetic Epidemiology Network of Arteriopathy” (phs001345) was performed at the Mayo Clinic Genotyping Core, and WGS was performed at the DNA Sequencing and Gene Analysis Center at the University of Washington (3R01HL055673-18S1) and the Broad Institute (HHSN268201500014C). We would like to thank the GENOA participants.

Genetic Epidemiology Network of Salt Sensitivity (GenSalt)

GenSalt utilizes a family feeding-study design. Each family is ascertained through a

proband with untreated prehypertension or stage-1 hypertension in rural China. Medical

history, lifestyle risk factors, and cold pressor tests are obtained at baseline visits while

BP, weight, blood and urine specimens are collected at baseline and follow-up visits^19^. The dietary intervention includes a 7-day low sodium-feeding (51.3 mmol/day), a 7-day high sodium-feeding (307.8 mmol/day), and a 7-day high sodium-feeding with an oral potassium supplementation (60 mmol/day). Whole genome sequencing (WGS) for the Trans-Omics in Precision Medicine (TOPMed) program was supported by the National Heart, Lung and Blood Institute (NHLBI). GenSalt was supported by research grants (U01HL072507, R01HL087263, and R01HL090682) from the National Heart, Lung and Blood Institute, National Institutes of Health, Bethesda, MD.

Genetics of Lipid Lowering Drugs and Diet Network (GOLDN)

GOLDN is a family-based study of European descent individuals recruited in Minneapolis and Salt Lake City (two of the NHLBI Family Heart Study sites). It aims to uncover genetic predictors of variability in lipid phenotypes, which include both fasting and postprandial lipids quantified using traditional methods, NMR, and high-throughput lipidomics. During the initial screening of ~1,350 individuals, the following criteria were used for exclusion: age < 18 years; fasting triglycerides ≥1500 mg/dL; recent history of myocardial infarction, coronary bypass surgery, or coronary angioplasty; self-report of a positive history of liver, kidney, pancreas, or gallbladder disease, or a history of nutrient malabsorption; current use of insulin; abnormal liver or kidney function; in women of childbearing potential, pregnancy, breastfeeding, not using an acceptable form of contraception. Of those who enrolled, 1,048 individuals consented to the use of their DNA in research; 893 participants with data on all exposures, outcomes, and covariates were included in the current study.

GOLDN biospecimens, baseline phenotype data, and intervention phenotype data were collected with funding from National Heart, Lung and Blood Institute (NHLBI) grant U01 HL072524. Whole-genome sequencing in GOLDN was funded by NHLBI grant R01 HL104135-04S1.

Whole genome sequencing (WGS) for the Trans-Omics in Precision Medicine (TOPMed) program was supported by the National Heart, Lung and Blood Institute (NHLBI). WGS for “NHLBI TOPMed: Genetics of Lipid Lowering Drugs and Diet Network” (phs001359) was performed at the University of Washington Northwest Genomics Center (3R01HL104135-04S1).

Hispanic Community Health Study - Study of Latinos (HCHS/SOL)

The Hispanic Community Health Study (HCHS)/Study of Latinos (SOL) is a multicenter,

community-based cohort study of Hispanic/Latino adults in the United States. The main

goal of the study is to identify risk factors which could either be protective or harmful to

the Hispanic community^20,21^. A total of 16,415 Hispanic/Latino individuals of age 18-74

were recruited. Participants are recruited in community areas surrounding four field

centers in the Bronx, Chicago, Miami, and San Diego. Whole genome sequencing

(WGS) for the Trans-Omics in Precision Medicine (TOPMed) program was supported by

the National Heart, Lung and Blood Institute (NHLBI). The Hispanic Community Health

Study/Study of Latinos was carried out as a collaborative study supported by contracts

from the National Heart, Lung, and Blood Institute (NHLBI) to the University of North

Carolina (N01-HC65233), University of Miami (N01-HC65234), Albert Einstein College

of Medicine (N01-HC65235), Northwestern University (N01-HC65236), and San Diego

State University (N01-HC65237).

Hypertension Genetic Epidemiology Network and Genetic Epidemiology Network

of Arteriopathy (HyperGEN)

The Hypertension Genetic Epidemiology Network Study (HyperGEN) - Genetics of Left

Ventricular (LV) Hypertrophy is a familial study aimed to understand genetic risk factors

for LV hypertrophy by conducting genetic studies of continuous traits from

echocardiography exams^22,23^. As part of HyperGEN study, four field centers recruited

African American and white hypertensive siblings, aged 23 to 87 years. Data from

detailed clinical exams as well as genotyping data for linkage studies, candidate gene

studies and GWAS have been collected and is shared between HyperGEN and the

ancillary HyperGEN - Genetics of LV Hypertrophy study. Whole genome sequencing

(WGS) for the Trans-Omics in Precision Medicine (TOPMed) program was supported by

the National Heart, Lung and Blood Institute (NHLBI). The HyperGEN Study is part of

the National Heart, Lung, and Blood Institute (NHLBI) Family Blood Pressure Program;

collection of the data represented here was supported by grants U01 HL054472 (MN

Lab), U01 HL054473 (DCC), U01 HL054495 (AL FC), and U01 HL054509 (NC FC). The

HyperGEN: Genetics of Left Ventricular Hypertrophy Study was supported by NHLBI

grant R01 HL055673 with whole-genome sequencing made possible by supplement -

18S1.

Jackson Heart Study (JHS)

The JHS is a large, population-based observational study evaluating the etiology of cardiovascular, renal, and respiratory diseases among African Americans residing in the three counties (Hinds, Madison, and Rankin) that make up the Jackson, Mississippi metropolitan area^24,25^. Data and biologic materials have been collected from 5,306 participants, including a nested family cohort of 1,498 members of 264 families. The age at enrollment for the unrelated cohort was 35-84 years; the family cohort included related individuals >21 years old. Participants provided an extensive medical and social history and had an array of physical and biochemical measurements and diagnostic procedures, and a subset of participants provided genomic DNA during a baseline examination (2000-2004) and two follow-up examinations (2005-2008 and 2009-2012), with a fourth examination ongoing. Annual follow-up interviews and cohort surveillance are ongoing.

Whole genome sequencing (WGS) for the Trans-Omics in Precision Medicine (TOPMed) program was supported by the National Heart, Lung and Blood Institute (NHLBI). WGS for “NHLBI TOPMed: The Jackson Heart Study” (phs000964.v1.p1) was performed at the University of Washington Northwest Genomics Center (HHSN268201100037C).

The Jackson Heart Study (JHS) is supported and conducted in collaboration with Jackson State University (HHSN268201800013I), Tougaloo College (HHSN268201800014I), the Mississippi State Department of Health (HHSN268201800015I/HHSN26800001) and the University of Mississippi Medical Center (HHSN268201800010I, HHSN268201800011I and HHSN268201800012I) contracts from the National Heart, Lung, and Blood Institute (NHLBI) and the National Institute for Minority Health and Health Disparities (NIMHD). The authors also wish to thank the staffs and participants of the JHS.

Multi-Ethnic Study of Atherosclerosis (MESA)

The Multi-Ethnic Study of Atherosclerosis is a National Heart, Lung and Blood Institute-sponsored, population-based investigation of subclinical cardiovascular disease and its progression^26^. A total of 6,814 individuals, aged 45 to 84 years, were recruited from six US communities (Baltimore City and County, MD; Chicago, IL; Forsyth County, NC; Los Angeles County, CA; New York, NY; and St. Paul, MN) between July 2000 and August 2002. Participants were excluded if they had physician-diagnosed cardiovascular disease prior to enrollment, including angina, myocardial infarction, heart failure, stroke or TIA, resuscitated cardiac arrest or a cardiovascular intervention (e.g., CABG, angioplasty, valve replacement, or pacemaker/defibrillator placement). Pre-specified recruitment plans identified four racial/ethnic groups (White European-American, African-American, Hispanic-American, and Chinese-American) for enrollment, with targeted oversampling of minority groups to enhance statistical power.

Whole genome sequencing (WGS) for the Trans-Omics in Precision Medicine (TOPMed) program was supported by the National Heart, Lung and Blood Institute (NHLBI). WGS for “NHLBI TOPMed: Multi-Ethnic Study of Atherosclerosis (MESA)” (phs001416.v3.p1) was performed at the Broad Institute of MIT and Harvard (3U54HG003067-13S1). Centralized read mapping and genotype calling, along with variant quality metrics and filtering were provided by the TOPMed Informatics Research Center (3R01HL-117626-02S1). Phenotype harmonization, data management, sample-identity QC, and general study coordination, were provided by the TOPMed Data Coordinating Center (3R01HL-120393-02S1), and TOPMed MESA Multi-Omics (HHSN2682015000031/HSN26800004). The MESA projects are conducted and supported by the National Heart, Lung, and Blood Institute (NHLBI) in collaboration with MESA investigators. Support for the Multi-Ethnic Study of Atherosclerosis (MESA) projects are conducted and supported by the National Heart, Lung, and Blood Institute (NHLBI) in collaboration with MESA investigators. Support for MESA is provided by contracts 75N92020D00001, HHSN268201500003I, N01-HC-95159, 75N92020D00005, N01-HC-95160, 75N92020D00002, N01-HC-95161, 75N92020D00003, N01-HC-95162, 75N92020D00006, N01-HC-95163, 75N92020D00004, N01-HC-95164, 75N92020D00007, N01-HC-95165, N01-HC-95166, N01-HC-95167, N01-HC-95168, N01-HC-95169, UL1-TR-000040, UL1-TR-001079, UL1-TR-001420, UL1TR001881, DK063491, and R01HL105756. The authors thank the other investigators, the staff, and the participants of the MESA study for their valuable contributions. A full list of participating MESA investigators and institutes can be found at <http://www.mesa-nhlbi.org>.

San Antonio Family Heart Study (SAFS)

The SAFHS began in 1991, and included 1,431 individuals in 42 extended families at baseline. Probands were 40 to 60 year old low-income Mexican Americans selected at random without regard to presence or absence of disease, almost exclusively from Mexican American census tracts in San Antonio, Texas. All first, second, and third degree relatives of the proband and of the proband’s spouse, aged 16 years or above, were eligible to participate in the study. As part of our ongoing studies, we have recruited new family members from the original families, expanding the cohort to almost 3,099 individuals primarily from 73 families. Our study is a mixed longitudinal design. Subjects have been seen between 1 and 4 times with an average of 1.95 examinations.

Whole genome sequencing (WGS) for the Trans-Omics in Precision Medicine (TOPMed) program was supported by the National Heart, Lung and Blood Institute (NHLBI). WGS for “NHLBI TOPMed: San Antonio Family Heart Study” (phs001215) was performed at the Illumina Genomic Services (3R01HL113323-03S1).

Collection of the San Antonio Family Study data was supported in part by National Institutes of Health (NIH) grants R01 HL045522, MH078143, MH078111 and MH083824; and whole genome sequencing of SAFS subjects was supported by U01 DK085524 and R01 HL113323. We are very grateful to the participants of the San Antonio Family Study for their continued involvement in our research programs.

Genome-wide Association Study of Adiposity in Samoans (Samoan)

The parent Samoan study is a population-based genome-wide association study (GWAS) of adiposity and cardiometabolic phenotypes among adults, 25-65 years of age, from the independent nation of Samoa in the South Pacific. The research goal of this study is to identify genetic variation that increases susceptibility to obesity and cardiometabolic phenotypes. Biomarker and questionnaire data were collected to assess cardiometabolic phenotypes. DNA was collected and the Affymetrix 6.0 chip used for SNP genotyping. After quality control checks on genotyping and excluding individuals with key missing data we have a final sample of 3,122 adults with high-quality genome-wide marker data^27^. Participation in TOPMed provided whole genome sequence data for 1,285 individuals from the GWAS sample chosen for maximal informativity for our Samoan-specific imputation panel.

Whole genome sequencing (WGS) for the Trans-Omics in Precision Medicine (TOPMed) program was supported by the National Heart, Lung and Blood Institute (NHLBI). WGS for “NHLBI TOPMed: Genome-wide Association Study of Adiposity in Samoans” (phs000972) was performed at the University of Washington Northwest Genomics Center (HHSN268201100037C) and the New York Genome Center (HHSN268201500016C).

Data collection was funded by NIH grant R01-HL093093 and R01-HL133040. We thank the Samoan participants of the study and local village authorities. We acknowledge the support of the Samoan Ministry of Health and the Samoa Bureau of Statistics for their support of this research.

Taiwan Study of Hypertension using Rare Variants (THRV)

The THRV-TOPMed study consists of three cohorts: The SAPPHIRe Family cohort,

TSGH (Tri-Service General Hospital, a hospital-based cohort), and TCVGH (Taichung

Veterans General Hospital, another hospital-based cohort), all based in Taiwan^28,29^.

1,271 subjects were previously recruited as part of the NHLBI-sponsored SAPPHIRe

Network (which is part of the Family Blood Pressure Program, FBPP). THRV is a

collaborative study between Washington University in St. Louis, LA BioMed at Harbor

UCLA, University of Texas in Houston, Taichung Veterans General Hospital, Taipei

Veterans General Hospital, Tri-Service General Hospital, National Health Research

Institutes, National Taiwan University, and Baylor University. THRV is based

(substantially) on the parent SAPPHIRe study, along with additional population-based

and hospital-based cohorts. Whole genome sequencing (WGS) for the Trans-Omics in

Precision Medicine (TOPMed) program was supported by the National Heart, Lung and

Blood Institute (NHLBI). The Rare Variants for Hypertension in Taiwan Chinese (THRV)

is supported by the National Heart, Lung, and Blood Institute (NHLBI) grant

(R01HL111249) and its participation in TOPMed is supported by an NHLBI supplement

(R01HL111249-04S1). SAPPHIRe was supported by NHLBI grants (U01HL54527,

U01HL54498) and Taiwan funds, and the other cohorts were supported by Taiwan

funds.

Women’s Health Initiative (WHI)

The Women’s Health Initiative (WHI) is a large study of postmenopausal women’s health investigating risk factors for cancer, CVD, age-related fractures and chronic disease^30^. It began in 1993 as a set of randomized controlled clinical trials (CT) and an observational study (OS). Specifically, the CT (n=68,132) included three overlapping components: The Hormone Therapy (HT) Trials (n=27,347), Dietary Modification (DM) Trial (n=48,835), and Calcium and Vitamin D (CaD) Trial (n=36,282). Eligible women could be randomized into as many as all three CTs components. Women who were ineligible or unwilling to join the CT were then invited to join the OS (n=93,676).

Whole genome sequencing (WGS) for the Trans-Omics in Precision Medicine (TOPMed) program was supported by the National Heart, Lung and Blood Institute (NHLBI). WGS for “NHLBI TOPMed: Women’s Health Initiative” (phs001237) was performed at the Broad Institute of MIT and Harvard (HHSN268201500014C).

The WHI program is funded by the National Heart, Lung, and Blood Institute, National Institutes of Health, U.S. Department of Health and Human Services through contracts 75N92021D00001, 75N92021D00002, 75N92021D00003, 75N92021D00004, 75N92021D00005. The authors thank the WHI investigators and staff for their dedication, and the study participants for making the program possible. A full listing of WHI investigators can be found at: <http://www.whi.org/researchers/Documents%20%20Write%20a%20Paper/WHI%20Investigator%20Long%20List.pdf>.

The content is solely the responsibility of the authors and does not necessarily represent the official views of the National Institutes of Health.

**References**

1.	Post, W. *et al.* Determinants of Coronary Artery and Aortic Calcification in the Old Order Amish. *Circulation* **115**, 717-724 (2007).

2.	Mitchell, B.D. *et al.* The genetic response to short-term interventions affecting cardiovascular function: Rationale and design of the Heredity and Phenotype Intervention (HAPI) Heart Study. *American Heart Journal* **155**, 823-828 (2008).

3.	Wright Jacqueline, D. *et al.* The ARIC (Atherosclerosis Risk In Communities) Study. *Journal of the American College of Cardiology* **77**, 2939-2959 (2021).

4.	Nadkarni, G.N. *et al.* Apolipoprotein L1 Variants and Blood Pressure Traits in African Americans. *Journal of the American College of Cardiology* **69**, 1564-1574 (2017).

5.	Hughes, G.H. *et al.* Recruitment in the Coronary Artery Disease Risk Development in Young Adults (Cardia) study. *Controlled Clinical Trials* **8**, 68-73 (1987).

6.	Redline, S., Schluchter, M.D., Larkin, E.K. & Tishler, P.V. Predictors of Longitudinal Change in Sleep-Disordered Breathing in a Nonclinic Population. *Sleep* **26**, 703-709 (2003).

7.	Larkin, E.K. *et al.* A Candidate Gene Study of Obstructive Sleep Apnea in European Americans and African Americans. *American Journal of Respiratory and Critical Care Medicine* **182**, 947-953 (2010).

8.	Cushman, M., Cornell, E.S., Howard, P.R., Bovill, E.G. & Tracy, R.P. Laboratory methods and quality assurance in the Cardiovascular Health Study. *Clinical Chemistry* **41**, 264-270 (1995).

9.	Fried, L.P. *et al.* The cardiovascular health study: Design and rationale. *Annals of Epidemiology* **1**, 263-276 (1991).

10.	Bowden, D.W. *et al.* Review of the Diabetes Heart Study (DHS) family of studies: a comprehensively examined sample for genetic and epidemiological studies of type 2 diabetes and its complications. *The review of diabetic studies: RDS* **7**, 188 (2010).

11.	Divers, J. *et al.* Genome-wide association study of coronary artery calcified atherosclerotic plaque in African Americans with type 2 diabetes. *BMC Genetics* **18**, 105 (2017).

12.	Kannel, W.B., Dawber, T.R., Kagan, A., Revotskie, N. & Stokes, J. Factors of Risk in the Development of Coronary Heart Disease—Six-Year Follow-up Experience. *Annals of Internal Medicine* **55**, 33-50 (1961).

13.	Kannel, W.B., Feinleib, M., McNamara, P.M., Garrison, R.J. & Castelli, W.P. AN INVESTIGATION OF CORONARY HEART DISEASE IN FAMILIES: THE FRAMINGHAM OFFSPRING STUDY. *American Journal of Epidemiology* **110**, 281-290 (1979).

14.	Splansky, G.L. *et al.* The Third Generation Cohort of the National Heart, Lung, and Blood Institute's Framingham Heart Study: Design, Recruitment, and Initial Examination. *American Journal of Epidemiology* **165**, 1328-1335 (2007).

15.	Vaidya, D. *et al.* Incidence of Coronary Artery Disease in Siblings of Patients With Premature Coronary Artery Disease: 10 Years of Follow-up. *American Journal of Cardiology* **100**, 1410-1415 (2007).

16.	Faraday, N. *et al.* Relation Between Atherosclerosis Risk Factors and Aspirin Resistance in a Primary Prevention Population. *American Journal of Cardiology* **98**, 774-779 (2006).

17.	Multi-Center Genetic Study of Hypertension. *Hypertension* **39**, 3-9 (2002).

18.	Daniels, P.R. *et al.* Familial aggregation of hypertension treatment and control in the genetic epidemiology network of arteriopathy (GENOA) study. *The American Journal of Medicine* **116**, 676-681 (2004).

19.	The GenSalt Collaborative Research, G. GenSalt: rationale, design, methods and baseline characteristics of study participants. *Journal of Human Hypertension* **21**, 639-646 (2007).

20.	LaVange, L.M. *et al.* Sample Design and Cohort Selection in the Hispanic Community Health Study/Study of Latinos. *Annals of Epidemiology* **20**, 642-649 (2010).

21.	Sorlie, P.D. *et al.* Design and Implementation of the Hispanic Community Health Study/Study of Latinos. *Annals of Epidemiology* **20**, 629-641 (2010).

22.	Arnett, D.K. *et al.* Sibling correlation of left ventricular mass and geometry in hypertensive African Americans and whites: The HyperGEN study. *American Journal of Hypertension* **14**, 1226-1230 (2001).

23.	Williams, R.R. *et al.* NHLBI Family Blood Pressure Program: Methodology and Recruitment in the HyperGEN Network. *Annals of Epidemiology* **10**, 389-400 (2000).

24.	Taylor, H.A. *et al.* Toward Resolution of Cardiovascular Health Disparities in African Americans: Design and Methods of the Jackson Heart Study. *Ethnicity & Disease* **15**, 4-17 (2005).

25.	Carpenter, M.A. *et al.* Laboratory, Reading Center, and Coordinating Center Data Management Methods in the Jackson Heart Study. *The American Journal of the Medical Sciences* **328**, 131-144 (2004).

26.	Bild, D.E. *et al.* Multi-Ethnic Study of Atherosclerosis: Objectives and Design. *American Journal of Epidemiology* **156**, 871-881 (2002).

27.	Minster, R.L. *et al.* A thrifty variant in CREBRF strongly influences body mass index in Samoans. *Nature Genetics* **48**, 1049-1054 (2016).

28.	Ranade, K. *et al.* A genome scan for hypertension susceptibility loci in populations of Chinese and Japanese origins*. *American Journal of Hypertension* **16**, 158-162 (2003).

29.	Wu, K.-D. *et al.* Clustering and Heritability of Insulin Resistance in Chinese and Japanese Hypertensive Families: a Stanford-Asian Pacific Program in Hypertension and Insulin Resistance Sibling Study. *Hypertension Research* **25**, 529-536 (2002).

30.	Design of the Women’s Health Initiative Clinical Trial and Observational Study. *Controlled Clinical Trials* **19**, 61-109 (1998).

**The Samoan Obesity, Lifestyle and Genetic Adaptations Study (OLaGA) Group**

Ranjan Deka, Dept. of Environmental Health, University of Cincinnati;

Nicola L. Hawley, Dept. of Chronic Disease Epidemiology, Yale University;

Stephen T. McGarvey, Dept. of Epidemiology and International Health Institute, and Dept. of Anthropology, Brown University;

Ryan L. Minster, Dept. of Human Genetics, University of Pittsburgh;

Take Naseri, Ministry of Health, Government of Samoa;

Muagututi‘a Sefuiva Reupena, Lutia I Puava Ae Mapu I Fagalele;

Daniel E. Weeks, Depts. of Human Genetics and Biostatistics, University of Pittsburgh.
